# Single-cell Sequencing Highlights Heterogeneity and Malignant Progression in Actinic Keratosis and Cutaneous Squamous Cell Carcinoma

**DOI:** 10.1101/2022.12.22.521622

**Authors:** Dan-Dan Zou, Ya-Zhou Sun, Xin-Jie Li, Wen-Juan Wu, Dan Xu, Yu-Tong He, Jue Qi, Ying Tu, Yang Tang, Yun-Hua Tu, Xiao-Li Wang, Xing Li, Feng-Yan Lu, Ling Huang, Heng Long, Li He, Xin Li

**Author notes:** Corresponding authors (L.H.), (X.L.). These authors contributed equally to this work (Dan-Dan Zou, Ya-Zhou Sun).

## Abstract

Cutaneous squamous cell carcinoma (cSCC) is the second most frequent of the keratinocyte-derived malignancies with actinic keratosis (AK) as a precancerous lesion. To comprehensively delineate the underlying mechanisms for the whole progression from normal skin to AK to invasive cSCC, we performed single-cell RNA-seq (scRNA-seq) to acquire the transcriptomes of 138,982 cells from 13 samples of six patients including AK, squamous cell carcinoma in situ (SCCIS), cSCC and their matched normal tissues, covering comprehensive clinical courses of cSCC. We identified diverse cell types, including important subtypes with different gene expression profiles and functions in major keratinocytes. In SCCIS, we discovered the malignant subtypes of basal cells with differential proliferative and migration potential. Differentially expressed genes (DEGs) analysis screened out multiple key driver genes including transcription factors (TFs) along AK to cSCC progression. Immunohistochemistry (IHC) / immunofluorescence (IF) experiments and single-cell ATAC sequencing (scATAC-seq) data verified the expression changes of these genes. The functional experiments confirmed the important roles of these genes in regulating cell proliferation, apoptosis, migration and invasion in cSCC tumor. Furthermore, we comprehensively described the tumor microenvironment (TME) landscape and potential keratinocyte-TME crosstalk in cSCC providing theoretical basis for immunotherapy. Together, our findings provide a valuable resource for deciphering the progression from AK to cSCC and identifying potential targets for anticancer treatment of cSCC.

## Introduction

Invasive cutaneous squamous cell carcinoma (cSCC) is the second most common skin malignancy accounting for 20% of keratinocyte carcinomas and the fatality rate is also second to melanoma [1]. The morbidity of cSCC is steadily increasing, posing a significant threat to public health. The most important cause of cSCC is ultraviolet (UV) irradiation from sunlight [2]. The occurrence of UV-induced cSCC is a multi-stage process, and its progression is usually slow [3]. Early detection, diagnosis and treatment are very important for patients with cSCC in the progressive multi-step process. The most significant risk factor for cSCC is actinic keratosis (AK), a precancerous lesion developed from the damage effects of chronical UV radiation, which has an extremely high incidence in the elderly. Up to 65% to 97% of cSCCs are reported to originate in lesions previously diagnosed as AKs [4]. The two diseases have a lot in common in terms of etiology, pathogenesis and genetic characteristics [5]. However, it is difficult to predict whether early precancerous lesions will further develop into invasive tumors [6]. Even though significant mutations of important genes closely related to cSCC were also detected in AK [7], most AK with these mutations did not transform into cSCC. Therefore, there is an urgent research need to define the critical molecular biomarkers and origin cancerous cells driving AK progress to cSCC, which will contribute to the prevention, early diagnosis, and effective treatment of cSCC.

At the same time, the occurrence, development, invasion and metastasis of tumors are closely related to the tumor microenvironment (TME) [8]. The TME includes immune and inflammatory cells, fibroblasts, microvessels and biomolecules infiltrated therein around tumor cells [9]. During the growth process, tumor cells interact with these cells and extracellular stroma, forming a special TME, affecting the production of chemokines, growth factors and proteolytic enzymes, and promoting tumor proliferation, angiogenesis, invasion and metastasis [10]. Recently, numerous studies have showed complex cellular communication network between tumor cells and TME in many types of cancer including cSCC [11]. Thus, the analysis of cell-cell communication in TME of cSCC will help us to understand the potential mechanisms during the progression from AK to cSCC in depth and develop new immunotherapy strategy for cSCC.

However, due to the complex tumor heterogeneity and high mutation load of cSCC [12], it is more difficult to identify the driving genes for the occurrence and development of cSCC. Although a number of cSCC related genes have been identified, the results in different studies vary greatly [7, 13]. In addition, due to limitations of technologies, the previous results based on bulk sequencing generally include a mixture of various cells, which may cover up key characteristic changes in tumor development [14]. Single-cell RNA sequencing (scRNA-seq) technology provides a powerful tool for obtaining transcriptome characteristics at the single-cell resolution level. It can not only reveal the heterogeneity of tumor cells and the progress of tumor development, but also reveal the intercellular communication between tumor cells and their TME [15]. Recently, single-cell sequencing technology has been applied in the studies of skin diseases, including skin aging, psoriasis and cSCC [16-18]. However, characterization of the initiation and progression of cSCC, especially the key transformation from AK to cSCC is still lacking.

In this study, we used scRNA-seq technology to analyze 138,982 cells from 13 samples of six patients including AK, squamous cell carcinoma in situ (SCCIS), cSCC and their matched normal tissues, covering comprehensive clinical courses of cSCC, filling the current blank of single-cell profiling of these diseases. Using this unique resource, we identified key cell subpopulations that may play an important role in the development from AK to cSCC. Importantly, we identified the early malignant cell subpopulation in SCCIS and comprehensively analyzed the characteristics related to the malignant status of these cells. Based on the identification of key cell subpopulations, we screened out key candidate genes of each important step in the transformation from normal skin to cSCC. The functional experiment verified that these key genes may play important driving roles in tumorigenesis. In addition, we described the TME landscape and cell-cell crosstalk of poorly-differentiated cSCC in details and identified important signaling pathways related to tumor progression. Together, our comprehensive analysis deeply revealed the whole malignant progression from normal skin to cSCC, and uncovered the heterogeneity of cSCC tumors, providing insights into understanding of cSCC initiation and progression and new therapeutic treatment development.

## Results

### Single-cell transcriptome profiling identified different subgroups of keratinocytes in human skin

We generated single-cell RNA-seq profiles of 13 samples from 6 patients presenting for surgical resection using the 10x Genomics Chromium platform. All these samples included 3 AK samples, 1 SCCIS tumor sample, 3 cSCC tumor samples (low-risk and high-risk) without any treatment and patient-matched 6 normal skin samples, which almost cover all clinical stages from AK to cSCC (Fig. 1, A and B; Table S1).

**Fig. 1.**
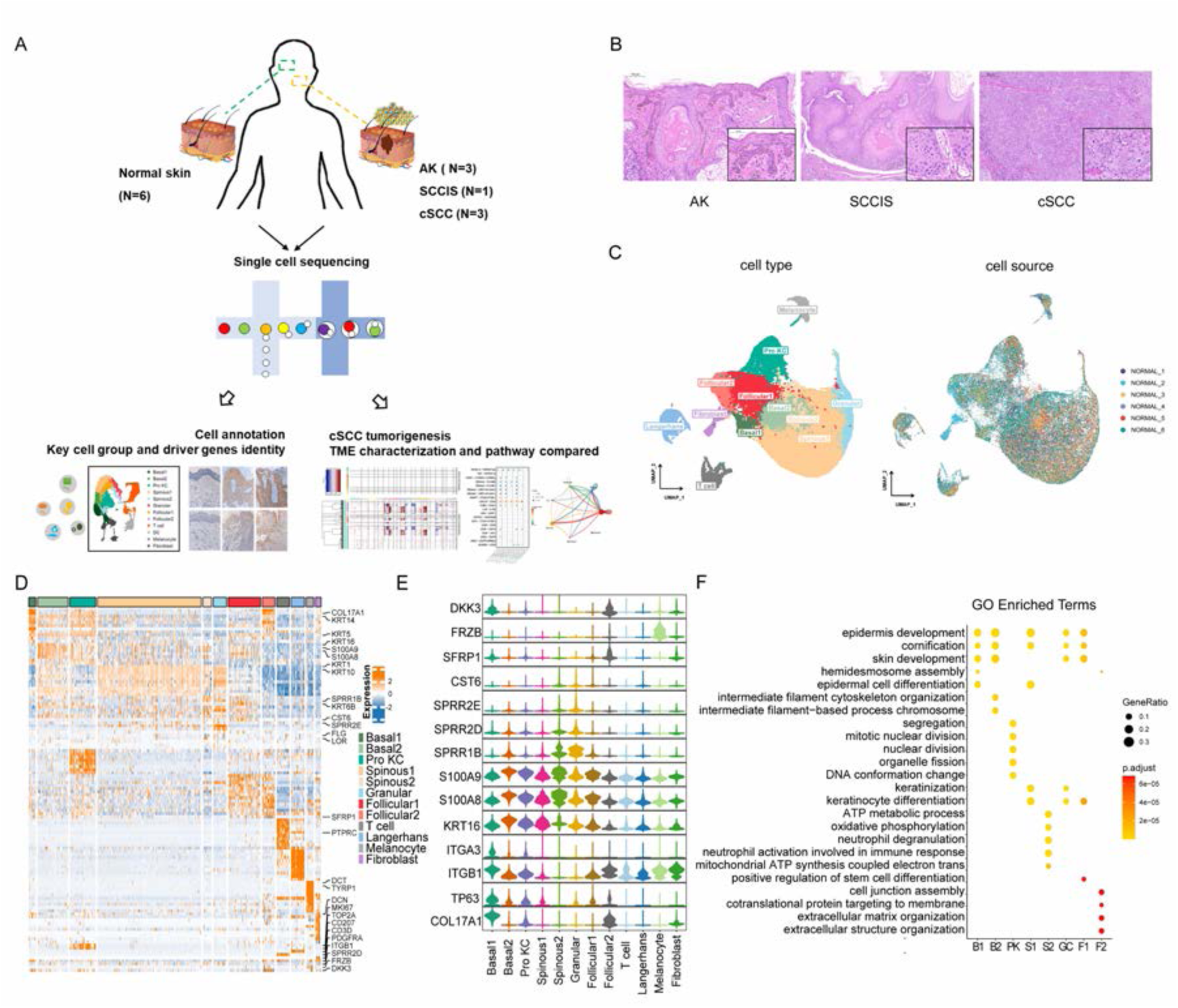
Single-cell transcriptome profiling identified different subgroups of keratinocytes in human normal skin. **(A)** Flowchart overview of single-cell sequencing in human skin of actinic keratosis (AK), squamous cell carcinoma in situ (SCCIS) and cutaneous squamous cell carcinoma (cSCC) patients. **(B)** Hematoxylin and eosin staining (H&E) of skin biopsies from representative AK (100X & 250X), SCCIS (50X & 250X) and cSCC (50X & 250X). **(C)** Uniform manifold approximation and projection (UMAP) plot of human normal skin labeled by cell type and patient respectively. **(D)** Heatmap showing gene expression signatures of each cell type. **(E)** Violin plot displaying the expression of representative genes to identify subpopulations for each cell type. **(F)** Representative gene ontology (GO) terms of signature genes in different cell subpopulations. The color keys from yellow to red indicate the range of p value.

We first explored the cellular composition of normal skin. After integration and initial quality control, we acquired single-cell transcriptomes in total of 57,610 cells from all six normal skin samples. Based on identified variably expressed genes across all normal skin cells, uniform manifold approximation and projection (UMAP) clustering identified 9 main clusters. Combining references with CellMarker [19], Panglao DB [20], Mouse Cell Atlas [21] and ImmGen [22] databases, we annotated each cell population based on their specific markers, including basal cells (COL17A1, KRT5, KRT14), spinous cells (KRT1, KRT10), granular cells (FLG, LOR), proliferating keratinocytes (Pro KCs) (MKI67, TOP2A), follicular cells (KRT6B, KRT17, SFRP1), Langerhans cells (CD207, CD1A), T cells (CD3D, PTPRC), melanocytes (PMEL, TYRP1) and fibroblasts (DCN, COL1A1) (Fig. 1, C and D).

Notably, we identified different subtypes of basal, spinous and follicular cells. UMAP analysis classified the keratinocytes into undifferentiated epidermal cells encompassing two subgroups of basal cells (Basal1 and Basal2) and Pro KCs, differentiated keratinocytes encompassing two spinous subpopulations (Spinous1 and Spinous2) and terminally differentiated cells (Granular) (Fig. 1C). Compared to Basal1, the expression levels of stemness markers (COL17A1, TP63, ITGB1, ITGA3) were decreased while inflammatory response genes (KRT16, S100A8, S100A9) were increased in Basal2 (Fig. 1, D and E). Functional gene ontology (GO) enrichment analysis of highly expressed genes of Basal1 and Basal2 subpopulations suggested that Basal1 were closely related to hemidesmosomes formation, while Basal2 were related to cell differentiation, migration and inflammatory response (Fig. 1F). Thus, we inferred that Basal1 were most likely the quiescent basal cells adhering to the basement membrane, which may represent epidermal stem cells. And Basal2 were cells that have finished proliferation to form the spinous layer for directional differentiation. In two subgroups of spinous cells, Spinous1 were associated with epidermal development, differentiation, and keratinization, while Spinous2 were associated with oxidative phosphorylation, neutrophil degranulation, and immune response (Fig. 1F). Compared with Spinous1, Spinous2 highly expressed small proline rich region proteins (SPRRs) encoding genes, such as SPRR1B, SPRR2D and SPRR2E (Fig. 1, D and E), which are involved in the formation of keratinocyte envelope [23]. Meanwhile, Spinous2 also highly expressed the cysteine protease inhibitor M/E (CST6), suggesting that Spinous2 subgroup was a well-differentiated upper spinous layer [24]. Follicular cells were also divided into two groups (Follicular1 and Follicular2). The functional enrichment of Follicular1 suggested that they were related to skin development and differentiation, while Follicular2 showed high levels of genes related to WNT signaling pathway (SFRP1, FRZB and DKK3) (Fig. 1, D and E). WNT signaling pathway plays a decisive role in regulating the functions of hair follicle stem cells, and inhibition of WNT signaling pathway can maintain the proliferation and inhibit the differentiation of stem cells [25]. Thus, the Follicular2 may represent outer bulge cells, which have been shown to secrete WNT inhibitors, influencing differentiation of the inner bulge.

In sum, we identified different subgroups in major types of keratinocytes including basal, spinous and follicular cells. The identification of these subgroups is important to understand the function and mechanisms of keratinocytes in human skin in depth and investigate the origin of cancer cells in the progression from normal skin to cSCC.

### Identification of potential key driver genes from normal skin to AK

To identify the potential drivers for AK, we first performed integration on all AK samples and patient- and site-matched normal samples. UMAP analysis of keratinocytes from AK and its corresponding normal skin samples showed that all AK and normal samples clustering were driven predominantly by cell type rather than patient. For all three AK samples in this study, the proportion of cell types of each sample was almost the same (Fig. 2A). These similar cell-type proportion suggested the low individual heterogeneity of AK samples.

**Fig. 2.**
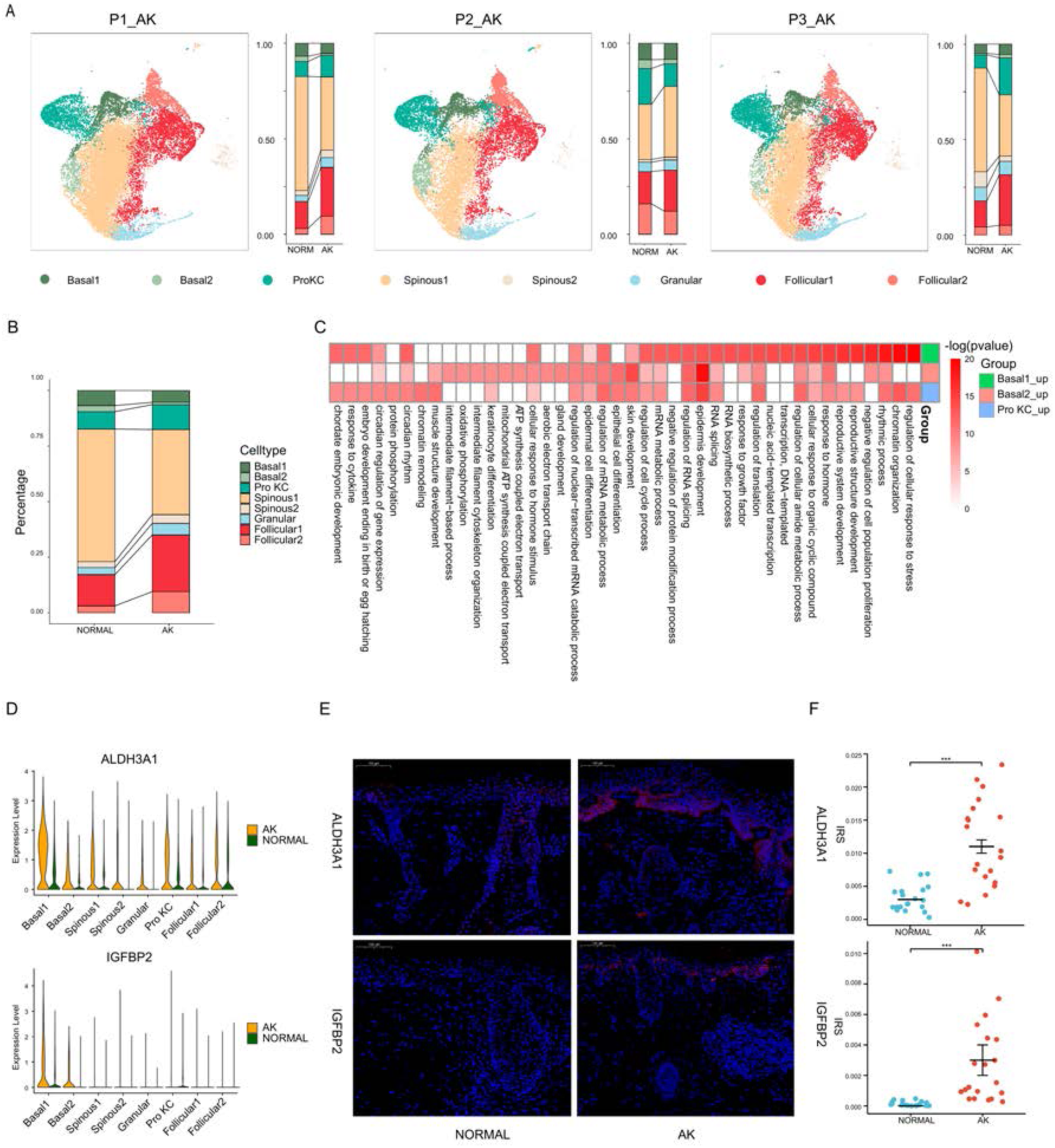
Identification of potential key genes driving normal skin to AK. **(A)** UMAP of scRNA-seq cells and cell proportion from each AK patient labeled by cell type. **(B)** Cell proportion of all AK samples and patient-matched normal skin samples. **(C)** Heatmap of GO terms for up-regulated genes in Basal1, Basal2 and Pro KC subpopulations for AK versus normal skin. **(D)** Violin plots showing the different expression levels of ALDH3A1 and IGFBP2 across cell types in AK and normal samples. **(E)** Left, immunofluorescence staining of ALDH3A1 and IGFBP2 in AK and normal skin groups. Scale bar, 100μm. Right, the mean optical density (IOD/Area) analyses of ALDH3A1 and IGFBP2 in AK and normal skin. n = 20 for each group. ***p < 0.001.

Compared to normal group, there was no significant difference in the proportion of basal cells and Pro KCs in AK group (Fig. 2B). This inferred that the proliferation and differentiation degree of keratinocytes in AK was not significantly different from that in normal samples. However, the proportion of spinous cells was slightly lower, and the proportion of follicular cells was slightly higher. It may be related to local epidermal atrophy in AK samples.

To further explore the mechanism of AK at the cell subpopulation level, we identified differentially expressed genes (DEGs) in major cell types of keratinocytes, especially cells with proliferation ability such as basal cells and Pro KCs between AK and normal. 549, 305 and 434 significantly up-regulated DEGs were identified in Basal1, Basal2 and Pro KCs subpopulations, respectively (Table S2-S4). GO enrichment analysis showed that it was mainly enriched in the terms associated with epidermal development, oxidative stress response, RNA metabolism, cell cycle, cytoskeleton, response to growth factors, etc. (Fig. 2C). An analysis between AK-related up-regulated DEGs and genes from the DisGeNET database [26] which collected genes and variants associated to human diseases revealed the high correlation of these genes and skin diseases such as dermatologic disorders, dermatitis, atopic, ichthyoses, acanthosis, etc. (Fig. S1A-C).

Combined with differential gene expression and functional enrichment analysis, we screened out a group of important candidate genes that may be closely related to AK occurrence and development (Fig. 2D and Fig. S1D, Table S5). Among the important candidate genes, some genes have been reported in previous studies showing close relationship with AK or related skin diseases. For example, CDKN2A is well known to take an important role in cSCC, and the latest study also found its mutation in AK [13, 27]. In our study, CDKN2A expression was increased in Basal1 and Basal2 subpopulations in AK stage, suggesting that CDKN2A may play a key role in the development of AK (Fig. S1D). For those genes that have not been reported, we selected seven candidate genes and verified their protein expression levels in an independent cohort including 20 pairs of facial AK and normal skin samples by immunofluorescence (IF). The results showed that the expression of ALDH3A1 and IGFBP2 was significantly upregulated in AK tissues and specifically mainly accumulated at the atypical keratinocytes of the epidermis (Fig. 2, E and F).

Acetaldehyde dehydrogenase 3A1 (ALDH3A1), as an important member of the acetaldehyde dehydrogenase superfamily, plays an important role in the occurrence and development of malignant tumors. DEG analysis showed ALDH3A1 was expressed in almost all keratinocytes and highly expressed especially in Basal1 cells of AK samples (Fig. 2D). Immunofluorescence experiment showed ALDH3A1 protein expression levels were significantly increased in 85% (17/20) AK tissues (Fig. 2, E and F). Previous study verified that ALDH3A1 could resist DNA damage caused by oxidative stress and genotoxicity in corneal epithelial cells to maintain epithelial homeostasis [28]. It suggested that ALDH3A1 might also participate in the DNA damage response in AK caused by UV irradiation. Recent study also found that ALDH3A1 could act as a prognostic biomarker and inhibit the epithelial mesenchymal transition (EMT) of oral squamous cell carcinoma (OSCC) through IL-6/STAT3 signaling pathway, further indicating its important role in SCC tumors [29].

Insulin-like growth factor binding protein 2 (IGFBP2), plays an important role in cell proliferation, differentiation, apoptosis and EMT. The high expression of IGFBP2 is significantly correlated with the malignant progression and prognosis of melanoma [30] and other tumors. In our study, IGFBP2 was specially highly expressed in Basal1 and Basal2 cells of AK samples (Fig. 2D). Immunofluorescence experiment showed IGFBP2 protein expression levels were significantly increased in 75% (15/20) AK tissues (Fig. 2, E and F). The up-regulation of IGFBP2 was also reported in both murine and human basal cell carcinoma (BCC), which promoted BCC development by mediating epidermal progenitor cell expansion via Hedgehog (Hh) signaling pathway [31]. It inferred that in AK development, IGFBP2 might also take an important role in promotion via related signaling pathways.

Collectively, these results indicated that from normal skin to AK, a lot of genes have changed expression. Especially, the basal cells specific up-regulated molecules ALDH3A1 and IGFBP2 likely contribute to the development of AK and may be the key driver genes in the process from photoaged skin to AK.

### Monotonically changed DEGs play important roles in the progression of AK to SCCIS

The individual P2 with both AK and SCCIS is a typical model to investigate the mechanisms of development from AK to SCCIS. We first integrated AK sample, SCCIS sample and normal adjacent skin sample from P2. As the epidermal parts were not separated from the dermal parts in SCCIS sample, it contained more non-keratinized cells including endothelial cells, T cells, vascular smooth muscle cells (VSMC), fibroblasts, etc. (Fig. 3A). The expression of cell proliferation and differentiation marker genes showed that the proportion of basal, Pro KC and terminally differentiated cells (Fig. 3A). Notably, the proportion of basal cells in SCCIS significantly increased compared with normal and AK samples, suggesting their specificity in SCCIS (Fig. 3B).

**Fig. 3.**
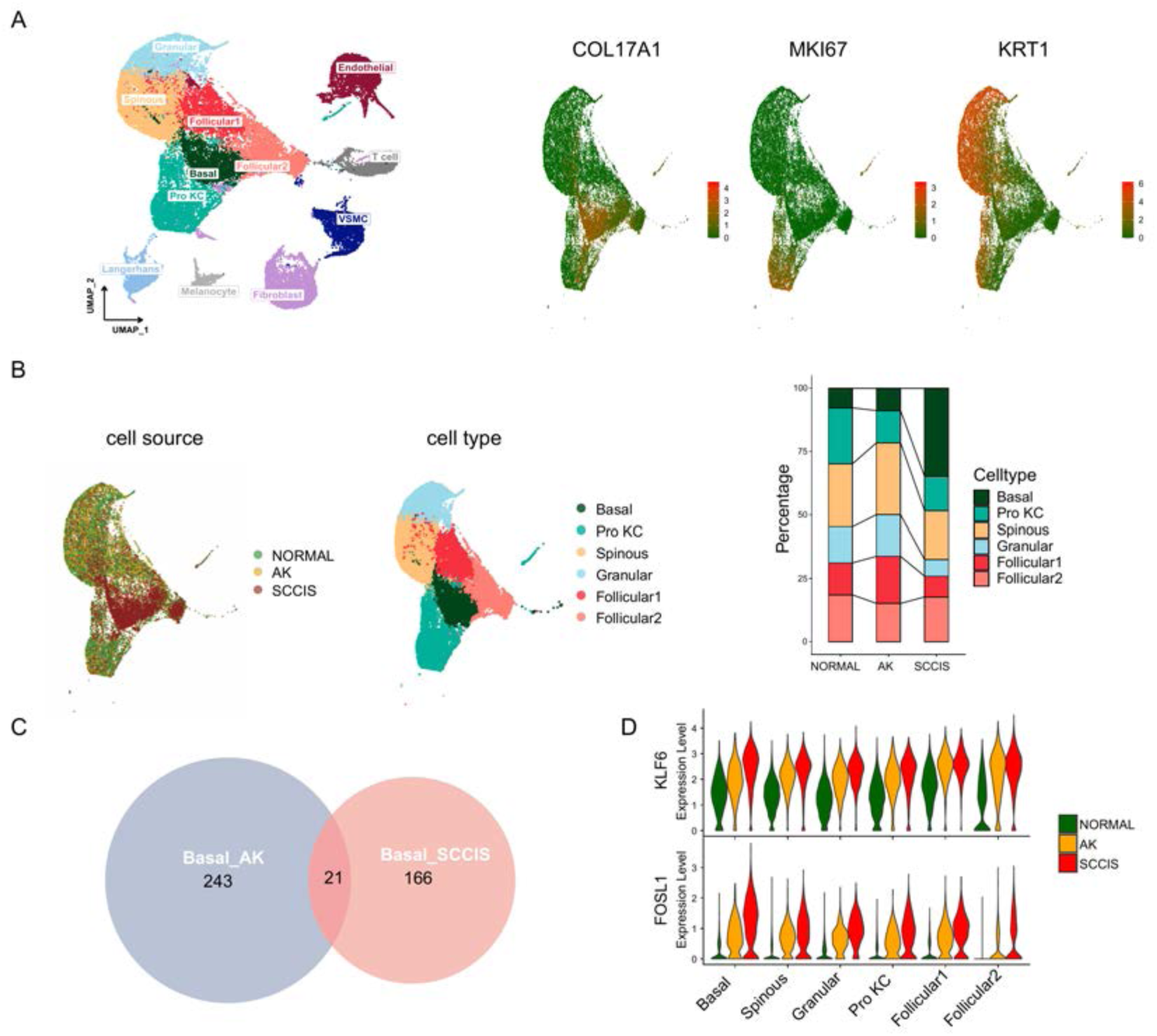
Monotonically changed DEGs play important roles in the progression of AK to SCCIS. **(A)** Left, UMAP of all cells from patient (P2) with both AK and SCCIS labeled by cell types; right, expression of basal, Pro KC and differentiated genes in all keratinocytes from P2. **(B)** Left, UMAP of all keratinocytes from P2 labeled by patient and cell type respectively; right, cell proportion of normal, AK and SCCIS samples in P2. **(C)** Overlap of up-regulated genes in basal cells from AK compared to normal and SCCIS compared to AK. **(D)** Violin plots showing the different expression levels of KLF6 and FOSL1 in normal, AK and SCCIS samples in P2.

To identify DEGs that monotonically increased during the process from normal skin to AK and SCCIS, which could be associated with the transformation from precancerous lesions to cancer, we first obtained AK up-regulated genes compared to normal, and SCCIS up-regulated genes compared to AK respectively in basal subpopulation, then got the overlap of these two gene sets. There are 21 overlapped up-regulated genes (Fig. 3C and Table S6), most of them were reported to have important functions in skin disease including cSCC. For example, the growth-controlling transcription factor Kruppel Like Factor 6 (KLF6) is an important contributor for epidermal decline and aging (Fig. 3D) [16]. The activator protein-1 (AP-1) family transcription factor subunit gene FOS Like 1 (FOSL1) is considered to be the potential driver of transformation from SCCIS to cSCC and is selectively highly expressed at the frontier of invasion in cSCC tumor cells (Fig. 3D) [32]. Another AP-1 family transcription factor subunit JunD Proto-Oncogene (JUND) can regulate cell proliferation, differentiation and apoptosis. It has been proposed to protect cells from p53-dependent senescence and apoptosis [33]. These data indicated that the constant up-regulation of these key growth-controlling transcription factors may play important roles in the process of AK to SCCIS. In addition, the ubiquitin binding protein Sequestosome 1 (SQSTM1) is involved in cell signal transduction, oxidative stress and autophagy. It was verified that SQSTM1 participated UV-induced decreased skin autophagy, and promoted the growth and progression of skin tumors through COX-2 [34]. The Ras Homolog Family Member B (RHOB) is a key regulator of UVB response. UVB-induced RHOB overexpression is involved in the initiation of cSCC by promoting the survival of keratinocytes with DNA damage mutations [35]. All of these findings suggested that these monotonically up-regulated genes are potential key drivers from precancerous lesion to carcinoma in situ during the development of cSCC, which may become potential targets for the prevention and treatment of cSCC.

In addition, we also investigated the monotonically down-regulated DEGs in this individual (Fig. S2A and Table S7). Although the number of these genes is also small, many of them showed strong associations with skin disorder or cancer, especially with cancer suppression. For instance, DNA Damage Inducible Transcript 4 (DDIT4) regulates apoptosis in response to DNA damage via its effect on mammalian target of rapamycin complex 1 (mTORC1) activity [36]. It is associated with skin atrophy [37]. The S100A family member S100A14 can regulate cell survival and apoptosis by modulating TP53 expression [38]. Levels of S100A14 have been found to be lower in cancerous tissue and associated with metastasis suggesting a tumor suppressor function [39]. Enolase 1 (ENO1) has been shown to bind to the c-myc promoter and function as a tumor suppressor [40]. Besides, the SPRR family member SPRR2A also showed constant downregulation. Recent study has identified it as a noninvasive biomarker in gastric cancer [41]. Considering it is the component of cornified keratinocyte cell envelope and a well-known keratinocyte terminal differentiation marker [42], we have reason to speculate the constant downregulation of SPRR2A indicating cancerization and increased invasiveness during the development from AK to SCCIS.

Although the underlying mechanisms of these genes in carcinogenesis need further functional studies, all the above data provided abundant evidences that these monotonically up-regulated and down-regulated DEGs act synergistically and play key driving roles in the formation of SCCIS from AK.

### Identification of malignant basal subpopulation in SCCIS

The increased proportion of basal cells in SCCIS hinted that they might be the crucial cell types in carcinomatous change of AK, thus we first investigated the characteristics of these cells. GO enrichment analysis showed that signature genes of basal cells in SCCIS were closely related to the biological processes of cell proliferation, morphological change, migration, cell connection and extracellular matrix, suggesting their invasive behavior (Fig. 4A). To define malignant cells, we calculated large-scale chromosomal copy umber variation (CNV) in each cell type of keratinocytes based on averaged expression patterns across intervals of the genome. We found that a subgroup of basal cells in SCCIS exhibited remarkably higher CNV levels than other basal cells (Fig. 4B). UMAP analysis of retrieved basal cells showed basal cells in SCCIS were divided into two major subgroups and basal cells with higher CNV levels were almost enriched in one subgroup (Fig. 4C). We further performed pseudotime analysis of these basal cells by selecting the subcluster with higher-expressed stem cell marker (COL17A1, TP63) as root cells. The result showed that basal cells of SCCIS differentiated into significantly distinct two subgroups, confirming that the distribution of the two subpopulations of basal cells was consistent with trajectory of differentiation (Fig. 4D and Fig. S2B).

**Fig. 4.**
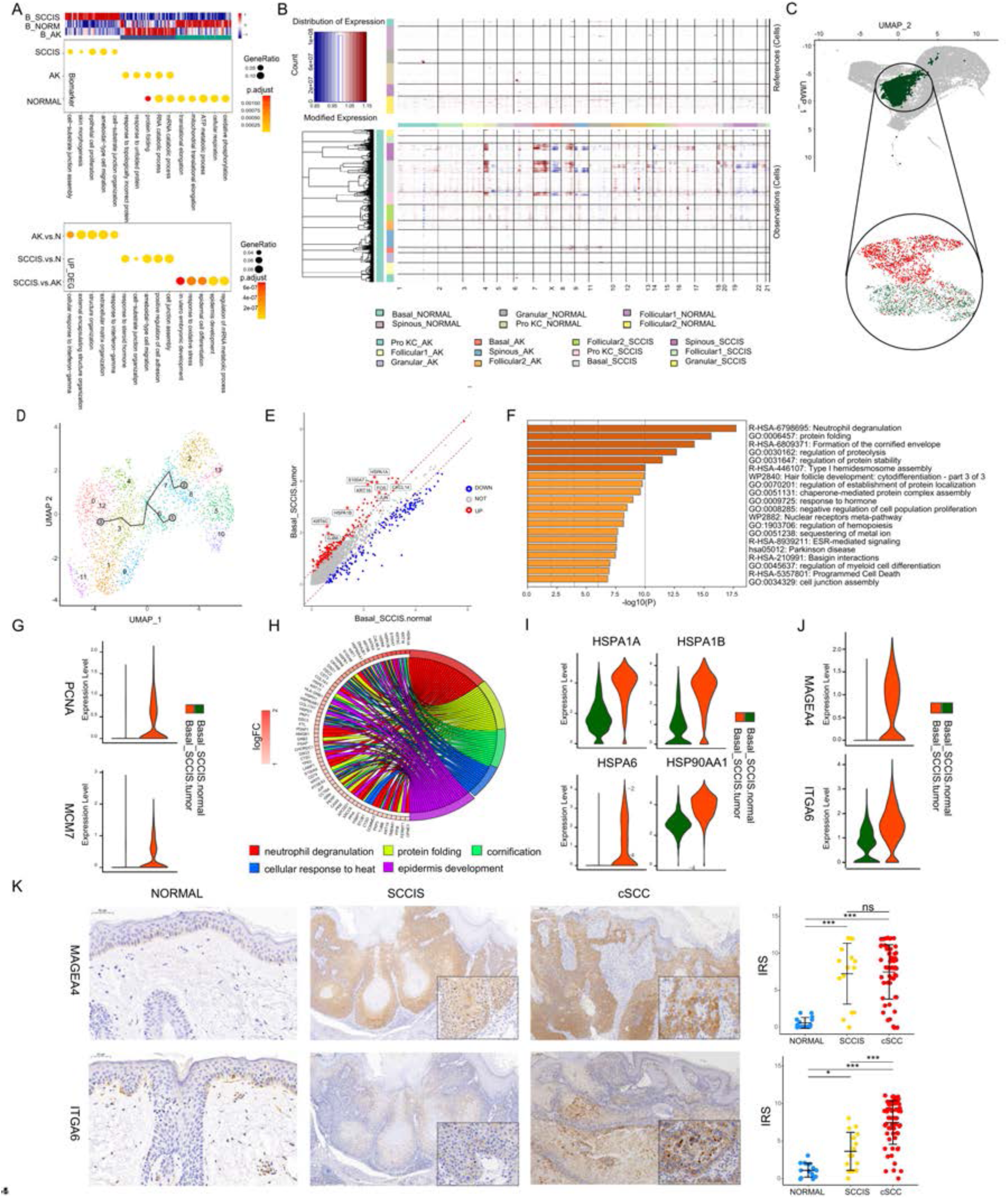
Identification of malignant basal subpopulation in SCCIS. **(A)** Representative GO terms for genes with specific expression in basal cell of SCCIS, AK and normal samples in P2 (upper) and up-regulated DEGs from AK versus normal, SCCIS versus normal, SCCIS versus AK respectively (lower). **(B)** Heatmap showing CNV levels of all keratinocytes from AK and SCCIS samples in P2. The keratinocytes from normal sample in P2 were defined as references. **(C)** UMAP of subgroups generated from basal cells in SCCIS sample showing basal cells with higher CNV level enriched in one subgroup; red dot representing basal cells with higher CNV level (cnv.score > 81.5). **(D)** Pseudotime analysis of basal cells in SCCIS was generated with Monocle3. **(E)** DEGs detected between Basal-SCCIS-tumor and Basal-SCCIS-normal. **(F)** Representative enriched Kyoto Encyclopedia of Genes and Genomes (KEGG) and GO terms in up-regulated genes (avg_log2FC > 0.58 and p_val_adj < 0.05). **(G)** Violin plots showing the expression level of representative DNA damage response marker genes in Basal-SCCIS-tumor and Basal-SCCIS-normal subgroups. **(H)** Chord plot showing the top up-regulated genes included in representative GO terms. **(I)** Violin plots showing the expression level of major members in HSP family across Basal-SCCIS-tumor and Basal-SCCIS-normal subgroups. **(J)** Violin plots showing the expression level of MAGEA4 and ITGA6 in Basal-SCCIS-tumor and Basal-SCCIS-normal subgroups. **(K)** Immunohistochemistry staining of MAGEA4 and ITGA6 in human skin of normal, SCCIS and cSCC samples.

The presence of these two different subpopulations of basal cells in SCCIS prompted us that they had different malignant status. We named the basal cells in SCCIS with higher CNV levels as Basal-SCCIS-tumor cells and the subgroup with lower CNV levels as Basal-SCCIS-normal cells. We next focused on the gene expression patterns in these two subpopulations and identified a total of 238 up-regulated genes in Basal-SCCIS-tumor cells (Fig. 4E and Table S8). GO and Kyoto Encyclopedia of Genes and Genomes (KEGG) enrichment analysis revealed that these genes were mainly associated with neutrophil degranulation, protein folding, keratosis, hemidesmosome assembly, cell proliferation, apoptotic signaling pathway, hemopoiesis, myeloid cell differentiation, stress response and cell junction organization (Fig. 4F). Notably, in Basal-SCCIS-tumor cells, DNA damage response related replication genes (PCNA, MCM7) were significantly up-regulated (Fig. 4G). Especially MCM7 is required for S-phase checkpoint activation upon UV-induced damage, which indicated the Basal-SCCIS-tumor cells were malignant cells from AK induced by UV-damage. In addition, a large number of heat shock protein (HSP) related genes (HSPA1A/B, HSP90AA1, HSPA6) were highly expressed in Basal-SCCIS-tumor cells, as well as activated keratin genes (KRT6A/B/C, KRT16, KRT17, KRT19) and S100 family genes (S100A7, S100A8, S100A9) (Fig. 4, H and I). HSPs play a role in tumor-related biological processes such as cell proliferation, apoptosis, invasion, tumor cell stemness, angiogenesis, glycolysis, hypoxia and inflammation. The HSP family is considered as a promising target for anticancer therapy. UV irradiation can induce keratinocyte injury and significant upregulation of heat shock proteins of *in vitro* skin model [43], which was consistent with our results (Fig. 4I). Previous studies have recognized that activated keratins are key early barrier alarmins, and the upregulation of these genes are involved in the alteration of proliferation, cell adhesion, migration, and inflammatory characteristics of keratocytes, leading to hyperproliferation and innate immune activation of keratocytes in response to epidermal barrier disruption [44]. The S100A family members were also reported to be significantly up-regulated in skin disorders or epithelial skin tumors. They participate in the immunoreactivity of keratocytes and have a potential utility as biomarkers for cancerous malignances [45]. Taken together, Basal-SCCIS-tumor cells with high CNV level may be highly invasive. As malignant cells, they may migrate and invade the dermis, and develop into invasive cSCC by promoting the proliferation ability of cells and the degradation of extracellular matrix to destroy the basement membrane.

### Basal-SCCIS-tumor specific genes were closely associated with progression from SCCIS to cSCC

Besides those genes were already reported closely related to cancerous malignances of SCCIS, we identified a group of candidate genes in the up-regulated genes in Basal-SCCIS-tumor subgroup that are closely related to tumor development. Combing with the single-cell transcriptomic data from invasive cSCC samples, we further screened out the candidate genes that were not only highly expressed in SCCIS samples, but also highly expressed in important keratinocytes in cSCC tumor samples, which play an important role in the progression of SCCIS to invasive cSCC (Fig. 4J and Fig. S2C, Table S9). These candidate genes were validated by immunohistochemistry (IHC) in an independent set of samples, including 15 normal skin tissues, 15 SCCIS tissues, and 60 invasive cSCC tissues (36 well-differentiated cSCC samples, 24 moderately-differentiated/poorly-differentiated cSCC samples), all of which were obtained from the faces of elderly patients.

Among them, MAGE family member A4 (MAGEA4) was found to be strongly positive in most SCCIS (73.33%) and invasive cSCC (76.67%). There was not significant difference of the expression between the well-differentiated cSCC group and moderately-differentiated/poorly-differentiated cSCC group (Fig. 4, K and I). MAGEA4 has been proven to inhibit p53-dependent apoptosis in cancer cells, enhance aggressivity of tumor cells, and induce cellular and humoral immune responses [46]. It was found to be highly expressed in melanoma, pancreatic cancer, lung cancer and esophageal squamous cell carcinoma [47]. Considering its potential utility as an indicator for malignancies of SCCIS tumor cells, we also performed immunofluorescence co-localization of COL17A1, PCNA and MAGEA4 in SCCIS tissues to investigate stemness and proliferative state of tumor cells. We found that in MAGEA4+ tumor cells of SCCIS, the stem cell marker COL17A1 and the proliferation marker PCNA were significantly increased compared with the adjacent tissues (Fig. S2D). However, there was significant individual heterogeneity in the expression of MAGEA4, and it was completely negatively expressed in some SCCIS and cSCC samples (Fig. 4, K and I). This inferred that MAGEA4 might become a promising new biomarker and target for the different subtypes of SCCIS with different invasive state. In addition, tumor-related gene Integrin Submit Alpha 6 (ITGA6) was also significantly overexpressed in Basal-SCCIS-tumor cells. The immunohistochemistry results showed that ITGA6 only scattered expression in the basal layer of the normal skin tissue and was up-regulated in SCCIS (P < 0.05) and invasive cSCC (P < 0.001), as well as being expressed at a higher level in invasive cSCC than in SCCIS (P < 0.001, Fig. 4K). We observed moderate to strong cytoplasmic and membranous positivity of ITGA6 in tumor cells. ITGA6 expression can indicate the progenitor potential of mesenchymal stem cells (MSC) [48]. Recent studies have reported that high ITGA6 expression enhances invasion and tumor-initiating cell activities in metastatic breast cancer (MBC) [49], providing evidence for the value of ITGA6 as cancer stem cell marker.

To further confirm the important role of MAGEA4 and ITGA6 in the development from SCCIS to cSCC, we performed functional experiment in human immortalized keratinocytes (HaCaT) and cSCC cell lines (A431, SCL-I, SCL-II). We first investigated the expression levels of MAGEA4 and ITGA6 in these cell lines and observed that the expression of MAGEA4 mRNA was extremely high in A431 cells, but not detectable in SCL-I and SCL-II cells (Fig. S3A), which is consistent with the immunohistochemical results observed in our clinical samples and Muehleisen et al [50]. We silenced the expression of MAGEA4 gene in A431 cells by siRNA (Fig. S3B), and the results showed that the proliferation, migration, invasive ability of A431 cells was significantly reduced (P < 0.01), while the apoptosis rate was increased (P < 0.01, Fig. S3C). The silencing of ITGA6 also significantly reduced the ability of proliferation, migration and invasion in the three cSCC cell lines (P < 0.01, Fig. S3, B, D, F and G), while apoptosis was significantly increased (P < 0.01, Fig. S3E). It is suggested that MAGEA4 and ITGA6 had a potential carcinogenic role in the progression of SCCIS to cSCC by regulating cell stemness, proliferation, apoptosis and extracellular matrix degradation.

### CNV scores positively correlated with malignant degrees of cSCC

To deeply investigate the genesis and key drivers of cSCC, we first integrated all three cSCC tumors and patient- and site-matched normal skin (Fig. 5A). These three individuals represent different malignant degrees of cSCC (Table S1). The cell-type proportion analysis and the expression of cell proliferation and differentiation marker genes reflected significant difference between tumor and normal samples (Fig. 5, B and C). There are more basal cells in tumor samples than in normal skin, which indicated the loss of terminal differentiation in tumor basal cells (Fig. 5, A-C). To define malignant cells, we identified large-scale CNV of keratinocytes based on averaged expression patterns across intervals of the genome. Keratinocytes in normal samples were set as reference cells. The presence of CNV in cSCC samples suggests that these cells may be tumor cells in squamous cell carcinoma samples. These tumor cells were mainly derived from Basal, Pro KCs, Follicular2 cells and a small number of Spinous cells (Fig. S4A). Then we compared the CNV landscapes among three patients. Poorly-differentiated cSCC individual exhibited remarkably higher CNV levels in most types of keratinocytes (Fig. S4B). In contrast, well-differentiated individual displayed low CNV scores (Fig. S4C), while moderately-differentiated individual had moderate CNV levels (Fig. S4D). This indicated that there was significant heterogeneity among different cSCC individuals and the CNV levels of individuals could reflect their malignant status.

**Fig. 5.**
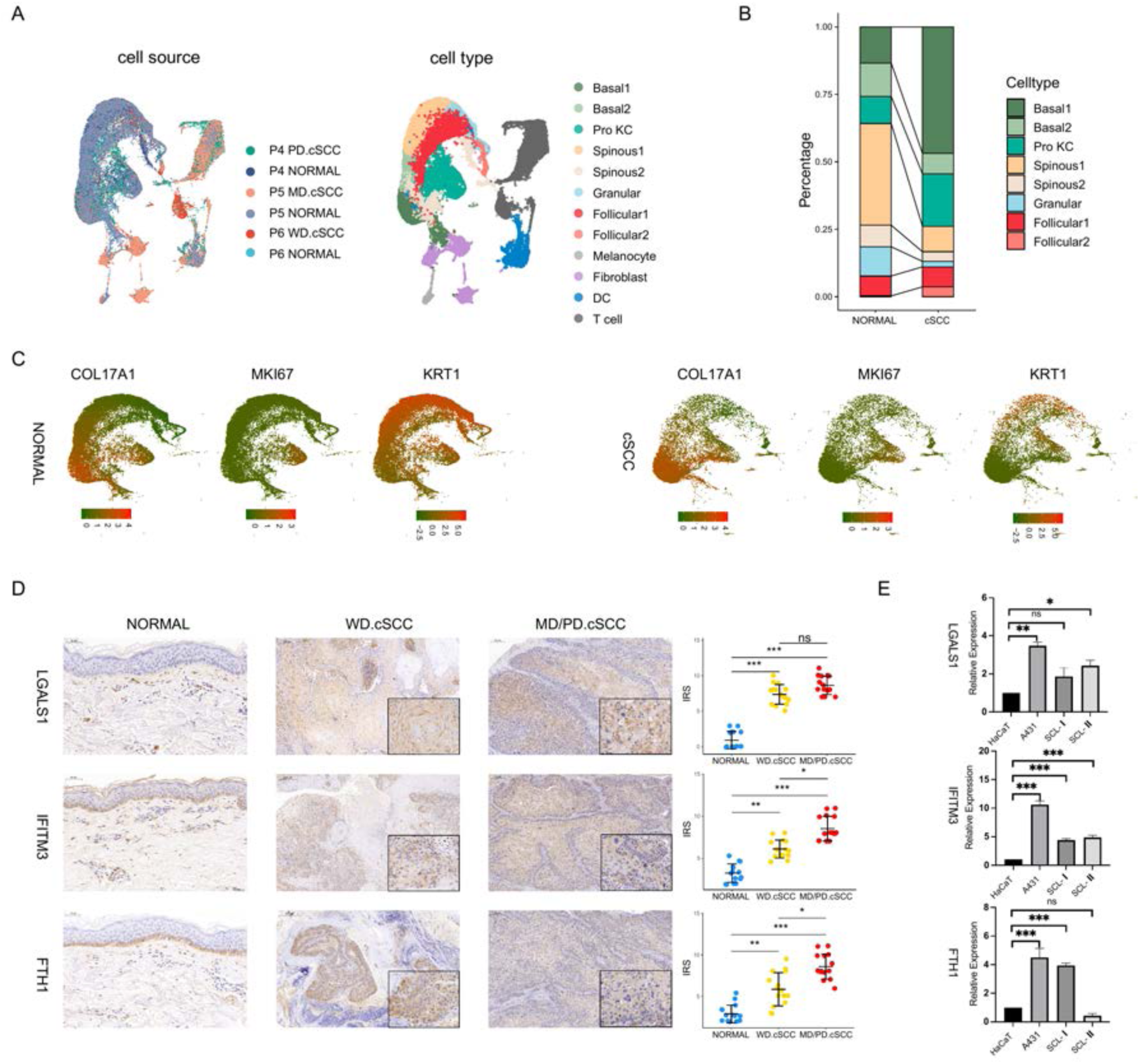
Identification of key genes associated with cSCC. **(A)** UMAP of all cells from cSCC patients labeled by sample and cell type respectively. **(B)** Cell proportion of keratinocytes in cSCC and normal groups. **(C)** Expression of basal, Pro KC and differentiated genes in all keratinocytes of cSCC and normal groups. **(D)** Left, immunohistochemical staining of LGALS1, IFITM3 and FTH1 in normal skin (200X), well-differentiated cSCC (WD.cSCC) (50X & 250X) and moderately-differentiated/poorly-differentiated cSCC (MD/PD.cSCC) (50X & 250X). Scale bar, 200 μm & 50 μm. Right, the immunoreactivity score (IRS) analyses of LGALS1, IFITM3 and FTH1 in normal skin, WD cSCC and MD/PD cSCC. n = 15 for each group. *p < 0.05; **p < 0.01; ***p < 0.001; ns, not significant. **(E)** The mRNA expression of LGALS1, IFITM3 and FTH1 in human immortalized keratinocytes (HaCaT) and cSCC cell lines (A431, SCL-I, SCL-II). *p < 0.05; **p < 0.01; ***p < 0.001; ns, not significant.

### Identification and functional characterization of key genes associated with cSCC

In order to understand the gene expression profile characteristics of invasive cSCC, we identified and analyzed gene function enrichment of significantly up-regulated DEGs in important cell subpopulations of cSCC compared to normal skin tissues. 778, 1044, 1159 and 760 significantly up-regulated DEGs were identified among Basal1, Basal2, Pro KCs, and Follicular2 cell subpopulations, respectively (Table S10-13). GO analysis was mainly concentrated in various tumor-related biological processes such as cell morphological change and adhesion, signaling pathway regulation, apoptosis and angiogenesis, as well as processes related to immunity including antigen processing and presentation, myeloid cell differentiation, and negative regulation of immune response, etc. (Fig. S5A). There were 888 and 247 significantly up-regulated DEGs in Spinous1 and Spinous2 cell subpopulations, respectively (Table S14-15). In addition to the biological process similar to basal cells, GO enrichment analysis showed that DEGs were mainly enriched in the regulation of programmed death, ATP metabolism and the production of type I interferon (Fig. S5A). Based on above differential gene expression and functional enrichment analysis, we identified a group of important candidate genes that may be closely related to tumor genesis and development in cSCC including CD74, CDKN2A, COL17A1, JUND, MMP1, BST2, LGALS1, IFITM3, ISG15, IFI6, FTH1, LAMA3, LAMC2, SAT1 and so on (Fig. S5B).

To verify the expression of those potential key genes with important functions in cSCC, we first performed immunohistochemistry experiment of these genes with significant differences in independent cohort including 30 cases of facial cSCC (15 well-differentiated cSCC samples and 15 moderately-differentiated/poorly-differentiated cSCC samples) and 15 cases of para-cancer normal skin tissues. Among them, 3 out of 8 genes were verified that they had significantly higher expression in cSCC group compared to normal group (Table S16). It was found that the protein expression levels of Galectin 1 (LGALS1), Interferon Induce Transmembrane Protein 3 (IFITM3) and Ferritin Heavy Chain 1 (FTH1) genes were significantly increased in cSCC (Fig. 5D). As a key promoter of angiogenesis and fibrosis, LGALS1 inhibits tumor immune response and is highly expressed in melanoma and head and neck cancer [51]. In this study, LGALS1 showed moderate to strong cytoplasmic and nuclear immunoreactivity in most well-differentiated and poorly-differentiated cSCC tumor cells, and some of them were weak staining, while para-cancer normal skin epidermal keratinocytes were almost negative. The LGALS1 expression in the tumor groups was significantly higher than that in the normal group (P < 0.001), but there was no statistical significance between the well-differentiated and poorly-differentiated cSCC groups (Fig. 5D). IFITM3 is an interferon-stimulating response related gene, which is related to cell proliferation, cell cycle regulation, autophagy, inflammation, EMT and many other processes. FTH1 is associated with iron metabolism, and may be involved in the protection of DNA from oxidative damage as well as in the regulation of inflammation, and tumor immune microenvironment. Although IFITM3 and FTH1 showed to moderate cytoplasmic/membranous immunoreactivity in basal cells in normal tissues, keratinocytes above basal showed almost negative expression. In poorly-differentiated cSCC, IFITM3 and FTH1 showed moderate to strong cytoplasmic, membranous immunoreactivity and a small number of nuclear staining. Medium to strong staining can also be seen at the leading edge or poor differentiated keratinocytes of the well-differentiated cSCC, while the expression is negative/weakly positive in the differentiated keratinocytes and the central keratinized areas of tumors (Fig. 5D). Overall, the expression of IFITM3 and FTH1 in poorly-differentiated cSCC was significantly higher than that of normal group (P < 0.001) and well-differentiated group (P < 0.05) (Fig. 5D). We also verified the mRNA expression levels of LGALS1, IFITM3, FTH1 in human immortalized keratinocytes (HaCaT) and human cSCC cell lines (A431, SCL-I, SCL-II). The results showed that these genes were significantly overexpressed in at least 2 human cSCC cell lines compared to HaCaT (Fig. 5E). Besides, although the Bone Marrow Stromal Cell Antigen 2 (BST2) and Spermine N1-Acetyltransferase 1 (SAT1) genes showed weak immunoreactivity in normal skin tissues and there were no statistically significant differences between cSCC and normal group (Fig. S6A), the mRNA expression levels of BST2 and SAT1 were significantly overexpressed in human cSCC cell lines compared to HaCaT (Fig. S6B).

To investigate the effects of LGALS1, IFITM3, FTH1, BST2 and SAT1 on the proliferation of human cSCC cells, the human cSCC cell lines A431, SCL-I, and SCL-II were transfected with small interfering RNA (siRNA) targeting these genes (Fig. 6A and Fig. S6C). The results showed that the silencing of LGALS1, IFITM3, FTH1, BST2 and SAT1 genes all inhibited the proliferation of the three human cSCC cells to varying degrees (Fig. 6B and Fig. S6D). These results suggested that LGALS1, IFITM3, FTH1, BST2 and SAT1 could regulate cell proliferation in cSCC. Then, Annexin V-FITC/propidium iodide (PI) staining and flow cytometry (FCM) was applied to quantify the effect of genes on apoptosis in human cSCC cells. The results showed that the gene silencing significantly increased the apoptosis rate of A431, SCL-I and SCL-II tumor cells (P < 0.01) (Fig. 6C and Fig. S6E). These results suggest that the up-regulation of LGALS1, IFITM3, FTH1, BST2 and SAT1 in cSCC may inhibit the apoptosis of tumor cells.

**Fig. 6.**
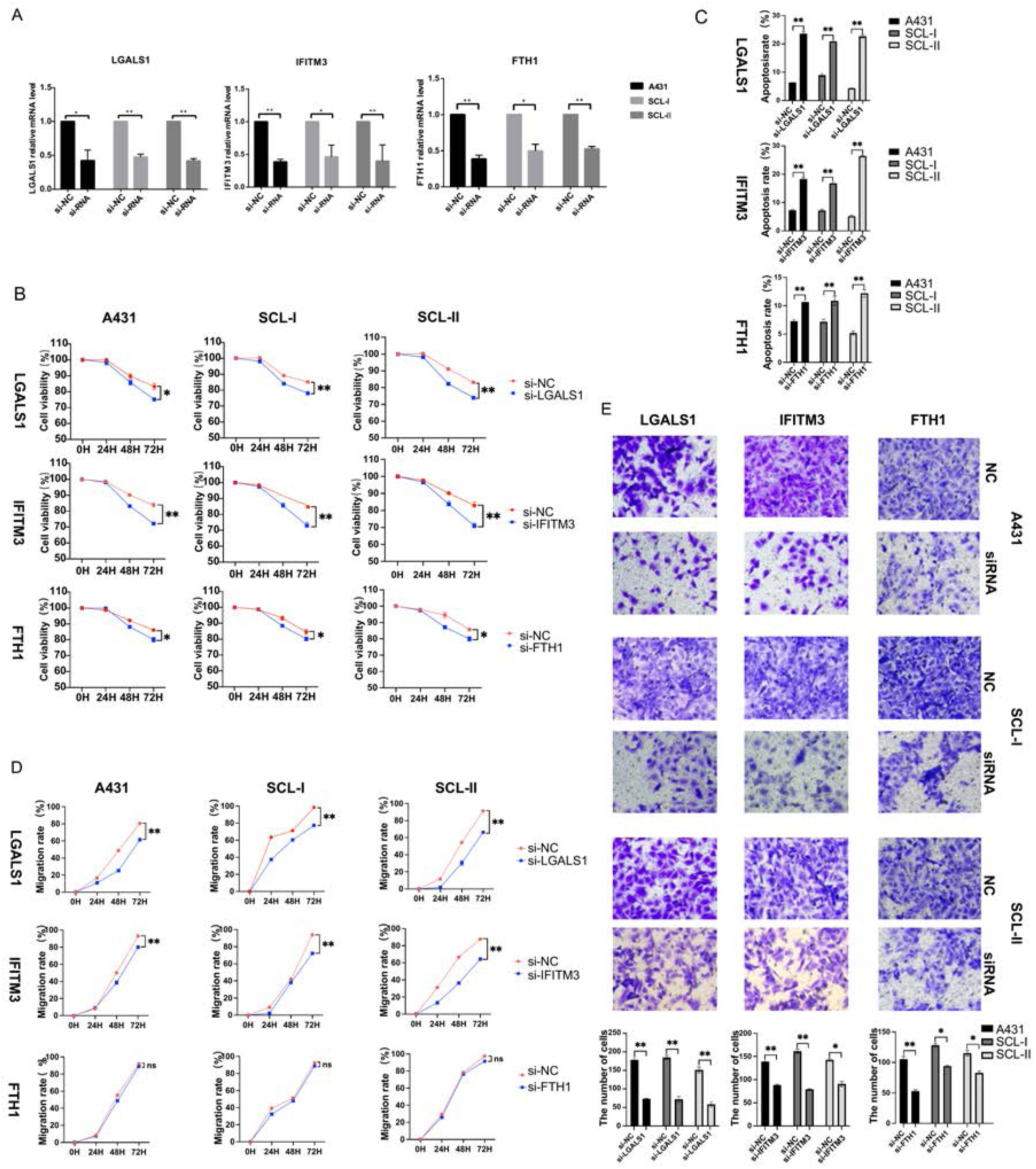
Functional characterization of key genes associated with cSCC. **(A)** Effect of siRNA on the expression of LGALS1, IFITM3 and FTH1 in A431, SCL-I and SCL-II determined by qRT-PCR. **(B)** Effect of LGALS1, IFITM3 and FTH1 on cSCC cell proliferation. The CCK-8 proliferation assay demonstrated a significant decrease in the proliferation of the si-LGALS1, si-IFITM3 and si-FTH1 groups compared with the si-NC group. *p < 0.05; **p < 0.01; ***p < 0.001; ns, not significant. **(C)** The effect of LGALS1, IFITM3 and FTH1 on cSCC cell apoptosis. Significant increase in the apoptosis of the si-LGALS1, si-IFITM3 and si-FTH1 groups compared with the si-NC group. *p < 0.05; **p < 0.01; ***p < 0.001; ns, not significant. **(D)** The scratch experiment showed that LGALS1 and IFITM3 knockdown resulted in a shorter vertical migration distance compared with the control group after 72 h, while there was no significant change in the si-FTH1 group. *p < 0.05; **p < 0.01; ***p < 0.001; ns, not significant. **(E)** Transwell assay showed that the invasion abilities of the si-LGALS1, si-IFITM3 and si-FTH1 groups significant decreased compared with the si-NC group. *p < 0.05; **p < 0.01; ***p < 0.001.

The effect of gene interference on the migration ability of human cSCC cells was detected by cell scratch assay. The results showed that after knocking down LGALS1, IFITM3, BST2 and SAT1, the migration distances of A431, SCL-I and SCL-II tumor cells were reduced after 72 h of scratching compared with the control group (P < 0.01), the tumor cells migration ability was decreased, while there was no significant difference with FTH1 in any tumor cells (P > 0.05) (Fig. 6D and Fig. S6F). These results suggest that up-regulation of LGALS1, IFITM3, BST2 and SAT1 may promote tumor cell migration ability in cSCC, whereas FTH1 has no great effect on this. Transwell invasion assay was used to detect the effect of gene interference on the invasion ability. The results showed that LGALS1, IFITM3, FTH1, BST2 and SAT1 gene silencing significantly reduced the invasion ability of A431, SCL-I and SCL-II (Fig. 6E and Fig. S6G). All these results suggested that LGALS1, IFITM3, FTH1, BST2 and SAT1 may take an important role in cSCC by regulating the processes of cell proliferation, apoptosis, migration and invasion.

### The tumor micro-environment (TME) landscape of cSCC

The progression of AK to cSCC is not only cancerization of keratinocytes, but also closely related to changes in skin microenvironment [10]. Long-term UVB irradiation can cause epidermal cell damage, while UVA can reach dermis and cause activation and oxidative damage of various cells in dermis [52]. For example, Langerhans may have disorders in cell number, migration ability, phenotypic changes and antigen presentation ability [52]. In addition, studies have confirmed fibroblast activation and expression of macrophage proteinases in the matrix, as well as loss of collagen XV and XVIII from the dermal basement membrane are early events in the progression of cSCC [10]. In order to understand the influence of tumor microenvironment on the occurrence and development of cSCC, we analyzed the non-keratinocytes lymphocytes, fibroblasts and dendritic cells and their cell communications in poorly-differentiated cSCC individual.

Tumor infiltrating lymphocytes are the main components of tumor microenvironment, especially T lymphocytes play an important role in immune response to tumor antigens. We identified eight lymphocyte subpopulations by re-clustering, including regulatory T cells (Treg), naive CD8+ T cells (CD8_Tnaive), CD8+ effector T cells (CD8_Teff), exhausted CD8+ T cells (CD8_Tex), CD4+ T cells (CD4_T), naive CD4 + T cells (CD4_Tnaive), CD4-CD8-naive T cells (DNT) and natural killer cells (NK) (Fig. 7A). Besides regulatory T cells, the exhausted CD8+ T subpopulation accounted for a considerable proportion. The exhausted CD8+ T cells have higher expression of inhibitory receptors, and the effector function is significantly reduced or lost, which may be one of the main factors of immune dysfunction. Ji et al. combined single-cell transcriptome with spatial transcriptome, and found that this group of cells were mainly located at the edge of inflammatory response and in immune cell clusters in cSCC [53].

**Fig. 7.**
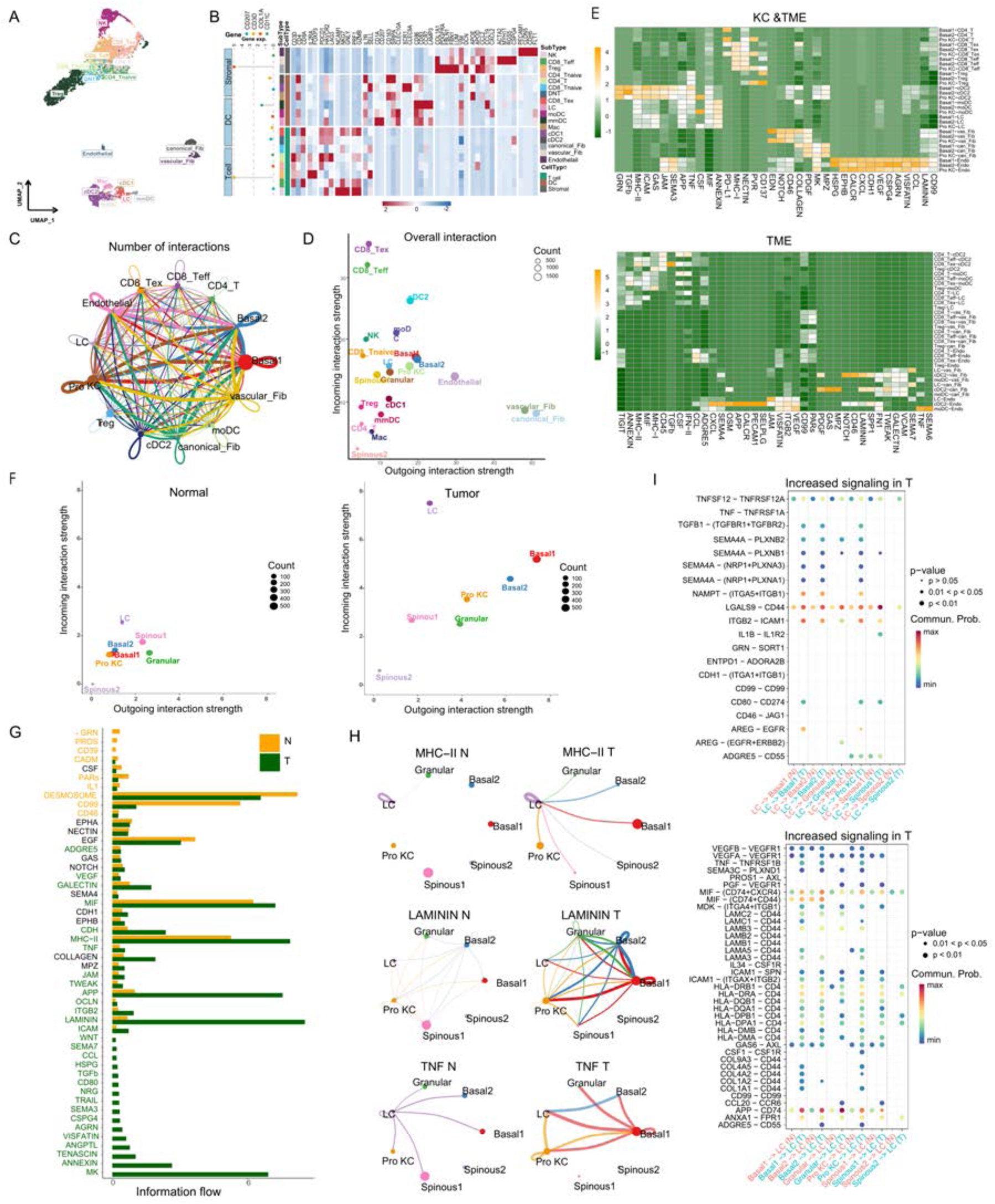
The analysis of cell-cell communication in TME of poorly-differentiated cSCC. **(A)** Identification of TME cell subpopulations, including T cells, DC cells and stromal cells. **(B)** Two-layered complex heatmap of selected cell marker genes in each cell cluster. Left, mean expression of known lineage markers; right, expression map of known marker genes associated with each cell subset. The relative expression values are scaled and transformed to a scale from -2 to 2. **(C)** Circle network diagram of overall cell–cell interactions. Thicker edge line indicates a stronger signal. **(D)** Comparison of total incoming path weights vs total outgoing path weights across cell populations. **(E)** Heatmap showing the communication probability on certain signaling pathway level. The top heatmap shows the cell-cell interactions between KC cells and TME cells. The bottom shows the interaction between subpopulations of TME cells. **(F)** Comparison of total incoming path weights vs total outgoing path weights between normal and tumor samples across common cell populations. **(G)** Significant signaling pathways were ranked based on differences in the overall information flow within the inferred networks between normal and tumor samples. The signaling pathways colored orange are enriched in normal tissue, and pathways colored dark green were enriched in the tumor tissue. **(H)** Circle plot showing the inferred intercellular communication signaling strength network between normal and tumor samples in MHC-II, LAMININ and TNF pathway. **(I)** Comparison of the significant ligand-receptor pairs between normal and tumor skin. The top shows the contribution to the signaling from Langerhans to KC subpopulations. The bottom shows the contribution to the signaling from KC subpopulations to Langerhans. Dot color reflects communication probabilities and dot size represents computed p-values. Empty space means the communication probability is zero. p-values are computed from one-sided permutation test.

Cancer-associated fibroblasts (CAFs) are one of the most important members of tumor microenvironment, interacting with tumor cells and playing an important role in the occurrence and development of tumor [54]. CAFs can secrete a variety of growth factors, cytokines and extracellular matrix proteins to promote tumor cell invasion and metastasis. In this study, we identified three subpopulations of CAFs including canonical_Fib, vascular_Fib and Endothelial (Fig. 7A). The expression levels of PDGFRA in canonical_Fib were higher than those in other subgroups (Fig. 7B). PDGFRA encodes platelet-derived growth factor receptor α, a cell surface receptor tyrosine kinase, which is activated by binding to the corresponding ligand PDGF and regulates cell division and proliferation. Abnormal gene activation of PDGFRA can lead to tumorigenesis and promote tumor angiogenesis, as well as induce macrophage migration to participate in immune regulation [55]. Vascular_Fib has a characteristic high expression of apolipoprotein APOE and COL18A1, which may be closely related to oxidative stress, inflammation and aging (Fig. 7B). Previous studies have found that tumor vascular endothelial cells can be transformed into CAFs under the regulation of TGF-β [56], and fibroblasts can also be transformed into vascular endothelial cells [57].

Dendritic cells (DCs) are important antigen presenting cells (APCs) in skin tissues, which play a key role in initiating, regulating and maintaining immune response [58]. DCs mainly contain Langerhans cells and dermal dendritic cells (DDCs). In general, epidermal Langerhans are immature, with weak antigen presentation capabilities. The mature DCs have enhanced antigen presentation capabilities, which can migrate to local lymph nodes to stimulate lymphocytes and activate immune responses [59]. Therefore, DCs play an important role in clearing skin tumor cells and preventing skin infections. In the tumor environment, the differentiation, development and maturation of DCs are interfered, which may help tumor cells evade immune surveillance[60]. In this study, six DC subpopulations were identified in poorly-differentiated cSCC individual (Fig. 7A). Among them, the stable monocyte derived DCs (Mo-DC) had characteristic high expression of CD14 and CLEC10A (Fig. 7B). The cluster with high expression of LAMP3 and CCR7 was identified as mature myeloid DCs (mmDC). It is a mature form of conventional dendritic cells, which has the potential to migrate from tumor to lymph node and can interact with a variety of T lymphocytes. Langerhans cells (LC) overexpressed CD1A, CD1C and CD207. In addition, we found that the expression level of IDO1 in Langerhans of poorly differentiated cSCC samples was significantly higher than that in normal skin samples (Fig. 7B). IDO1 is a classical tolerogenic mediator which not only engenders immune tolerance to tumor antigens, inhibits T cell cytotoxic activity and promotes differentiation into Tregs, but also acts in pathogenic inflammatory processes, may be a potential target for the development of oncology therapeutic inhibitors [61]. The immature conventional type I dendritic cells (cDC1) expressed unique C-type lectin receptor CLEC9A and chemokine receptor XCR (Fig. 7B). They can cross-present antigen and promote anti-tumor immune response of CD8+ T cells. Meanwhile, type II dendritic cells (cDC2) expressed CD163 and SIRPA. Another cluster highly expressed CD68, and represent macrophages (Mac) (Fig. 7B).

### Cell-cell communication analysis revealed important signaling pathways related in cSCC tumor

To investigate the effect of TME on invasive cSCC, CellChat, a cell communication analysis tool for single cell transcriptome data, was used to analyze the intercellular interactions of cSCC samples based on gene expression data, ligand-receptor database information and cell communication reference database (CellChatDB) [62]. It was found that cell-to-cell interaction was significantly enhanced in poorly-differentiated cSCC, and multiple interaction pathways were significantly active. The interactions between Basal1, Basal2 and Pro KCs and other cells were significantly increased, suggesting that these cell populations were the most important participants in cell crosstalk with TME during the development of cSCC (Fig. 7, C-F). In non-keratinocytes, fibroblasts released the most signal intensity, while CD8+ effector T cells received the most signal, canonical_Fib was particularly active in interaction with other cells (Fig. 7, D and E).

Further analysis revealed that some classical cancer-related signaling pathways were significantly altered in invasive cSCC (Fig. 7G). Basal cells can interact with various immune cells through Laminin and TNF signaling pathways (Fig. 7, E, G-I). Laminin is an important component of extracellular matrix (ECM), which is involved in basement membrane skeleton formation, cell adhesion, growth, differentiation and migration [63]. Laminin signaling pathways have been proven to regulate the morphology, differentiation and movement of a variety of cells including keratinocytes [64]. They also can participate in signal transmission and promote tumor infiltration and metastasis [65]. In cSCC, the expressions of ligand-receptors corresponding to Laminin signaling pathway were significantly enhanced in keratinocytes, suggesting that laminin signaling pathway has important significance for the occurrence and development of cSCC. The TNF family and its receptors play key roles in a variety of immune and inflammatory process. It plays a dual role in tumor, not only playing an immunomodulatory and tumor suppressive role, but also promoting tumor immune escape by inducing inflammatory response, promoting tumor cell survival, proliferation and EMT, regulating Treg and bone marrow derived suppressor cells (MDSCs) [66]. In addition, cell interaction between Basal cells, Pro KCs and Langerhans cells was enhanced through MHC-II and ICAM signaling pathway (Fig. 7, E, G-I). Studies have found that some tumor cells can play the role of antigen presentation by upregulation of MHC-II expression on cell surface, inducing differentiation and invasion of Tregs cells and participating in tumor genesis [67]. ICAM-I and its ligands LFA-1, MAC-1 and CD18 interact to regulate antigen presentation and migration or adhesion of various inflammatory cells [68]. These results suggested that these cell subpopulations and related signaling pathways may play key roles in cSCC.

### Chromatin accessibility is associated with transcription factor activity

To understand the chromatin accessibility and its role on regulation of gene expression in invasive cSCC, we also performed scATAC-seq on tumor sample of poorly-differentiated individual with scRNA-seq to generate paired, cell-type specific chromatin accessibility and transcriptional profiles. We leveraged the annotated scRNA-seq dataset of tumor sample to predict scATAC-seq cell types with Seurat using label transfer. Comparison between scATAC-seq cell-type predictions and curated annotations of scRNA-seq dataset indicated that all major cell types were present in both datasets (Fig. S7A). Then we detected accessible chromatin regions and investigated differentially accessible chromatin regions (DARs) between cell types (Table S17). The majority of DARs were located in a promoter region within 1 kb of the nearest transcriptional start site, especially in Basal1 the proportion was more than 95% (Fig. S7B). Meanwhile, a lot of DARs were closely associated with DEGs in their respective cell types and we could distinguish cell groups in scATAC-seq dataset based on the state of DARs. For example, the ATAC peaks of KRT5 were increased in major keratinocytes. CD83 is one of the best markers for mature DCs [69]. The coverage plot showed an increase in number and amplitude of ATAC peaks within its promoter and gene body in DCs and Langerhans (Fig. S7C).

Given that many transcription factors may be key determinants in the development of cSCC in our above results, we used chromVAR to infer transcription-factor associated chromatin accessibility in scATAC-seq dataset of cSCC tumor. We observed that the cell types also could be distinguished by transcription factor activities, suggesting these cell-type-specific transcription factors could regulate chromatin accessibility. For example, as well-known driver genes in cSCC, TP63 and TP53 were detected an enrichment of their binding motifs within DAR in major keratinocytes (Fig. S7D and Table S18) that was supported by increased chromatin accessibility and increased transcription in the scRNA-seq of them. More importantly, similar patterns were seen for FOSL1, which were identified as potential key driver transcription factors for the development of cSCC in our above results (Fig. S7D and Table S18). In addition, predicted cis-regulatory chromatin interactions by Cicero also showed a positive correlation between transcription factor activity and expression on a global level (Fig. S7E). Meanwhile, the above key transcription factors including TP63 and FOSL1 showed a positive correlation between motif activity and expression, further supporting their important driving roles in the development of cSCC as transcriptional activators (Fig. S7F and Table S19).

## Discussion

The occurrence and development of tumor is an extremely complex process. Although many studies have been carried out, the key mechanisms of the occurrence and development of AK and cSCC are still elusive. In this study, the gene expression profile information from 13 samples of 6 patients was obtained by single-cell transcriptomic sequencing, covering all stages of cSCC development including normal skin, AK, SCCIS and invasive cSCC, and 138,982 high-quality single cells were finally obtained for analysis.

The six normal skin tissues near the lesions were all from the exposed sites of elderly individuals, showing obvious photoaging, but there were no obvious pathological changes observed by visual inspection neither morphological abnormality of keratinocytes observed by histopathology. Nine different main cell clusters were identified from normal skin tissues and further subgroup analysis found different subtypes in basal, spinous and follicular cells. There have been reports about the subgroups of epidermal cells in previous studies [17, 18]. In our study, according to the expression of marker genes, Basal1 subgroup had high expression of the main components of hemidesmosomes COL17A1 and stem cell marker genes, suggesting that Basal1 might be stationary Basal cells attached to the basement membrane [70]. Although Basal2 subgroup had high expression of basal cell-related markers, the expression of COL17A1 and stem cell-related genes was decreased, suggesting they may be the basal cells that have finished division and are about to directionally differentiate. The Pro KCs highly expressed MKI67 and also had basal cell markers. They are transient amplifying cells (TAs) with strong proliferating ability, which can leave from the basal layer after limited mitosis and enter the process of terminal differentiation and migration [71]. Follicular cells can also be divided into Follicular1 and Follicular2 subgroups. Follicular2 had higher expression of WNT pathway related suppressor genes such as SFRP1, FRZB, and DKK3 than that of Follicular1. Cheng et al. also identified these cells in human normal skin by single-cell sequencing technology and speculated that these cells were stem cell groups in the protuberant of the outer root sheath of hair follicles [18, 25].

Our results showed that the percentage of Basal cells in SCCIS increased significantly compared with AK and normal samples. This was even more significant in cSCC, especially in the Basal1 subpopulation with high expression of stem cell-related markers, while Pro KCs only slightly increased. These results suggested that cell terminal differentiation may be impaired during the development of cSCC, and that basal cells play a more important role than Pro KCs.

Although the origin of cSCC has always been controversial, it is believed that basal cells with rapid proliferation ability and differentiated keratinocytes may all be the origin cells of cSCC [72]. However, there are other studies suggest that stem cells such as static epidermal stem cells and hair follicle stem cells are the most important origin cells of cSCC [73]. Morris et al. found that skin tumors came from stationary, 5-Fluorouracil-insensitive epidermal stem cells rather than rapidly proliferating epidermal cells in mouse chemocarcinogenic model of cSCC [74]. Adriana et al. also found that epidermal stem cells were the main origin cells of basal cell carcinoma, and proliferating epidermal cells mainly caused benign proliferative skin lesions [75].

To identify the malignant keratinocytes in AK, SCCIS and cSCC, we performed CNV analysis. Significant CNV differences were identified in cSCC samples, and the CNV differences among the samples were significant, which was proportional to the histopathological classification and risk grade, confirming the significant heterogeneity among the cSCC samples. However, we did not identify obvious CNV in AK samples, which may be related to the low proportion of malignant cells or the mild degree of malignancy in AK samples. In SCCIS samples, we also identified some cells with CNV differences, which may be the keratinocytes with early malignant transformation. Identification of malignant cells from SCCIS and invasive cSCC showed that these cells were mainly derived from stationary basal cells, some Pro KCs, Follicular2 cells and a small number of spinous cells, again suggesting the importance of Basal cells. Therefore, we focused on the characterization of these cells in SCCIS to understand the key events that promote the progression of precancerous lesions or carcinoma in situ to invasive cSCC. The characteristic marker genes of Basal cells of SCCIS were compared with normal and AK Basal cells. Two subpopulations of Basal cells in SCCIS were identified based on CNV scores and reclustering and Basal-SCCIS-tumor with higher CNV score were identified as malignant cell group.

Ji et al. identified a group of tumor-specific keratinocyte (TSK) subgroups in European cSCCs by using scRNA-seq and revealed their spatial distribution in combination with spatial transcriptomics [53]. In this study, we did not identify identical TSK cell subpopulations, which may be related to ethnic differences, different sample sources, and significant heterogeneity between tumors. However, we found a unique subpopulation in early stage of cancer with certain invasive characteristics in SCCIS and successfully identified malignant cells with significant CNV in invasive cSCC, revealing the heterogeneity of keratinocytes in different stages of AK and cSCC.

We compared the expression profiles of the same cell subpopulations during progression of AK, SCCIS and cSCC and identified a group of important candidate genes in each disease stage. ALDH3A1, IGFBP2, DYNC1H1, NFKBIZ and RND3 were specifically highly expressed in AK. ALDH3A1 is a newly discovered tumor stem cell marker in recent years. Overexpression of ALDH3A1 in melanoma and lung cancer not only regulates tumor cell stemness and the process of EMT, but also promotes inflammation through up-regulation of inflammatory factors such as COX2 and PGE2, and enhances the expression of PD-L1 to affect immune escape [76]. In vitro studies have confirmed that IGFBP2 is involved in regulating the proliferation, invasion and metastasis of tumor cells. IGFBP2 secreted in melanoma activates the PI3K/Akt pathway to promote tumor angiogenesis by binding to integrin αVβ3 [77]. However, the expression of ALDH3A1 and IGFBP2 was not elevated in the three cSCC samples, suggesting that they may play different regulatory roles in different stages of AK and cSCC.

It is of concern that two basal subgroups with different levels of CNV were identified in SCCIS, and the expression of a large number of heat shock proteins was generally increased in the Basal-SCCIS-tumor subgroup with higher level of CNV. Fernandez et al. found that HSP70 was increased in the cytoplasm of keratinocytes in cSCC tissues arising from AK and was positively correlated with dermal infiltration level [78]. It may be an early potential marker of progression from AK to cSCC. The high expression of activated keratin genes and S100 family genes also indicated the high invasiveness of Basal-SCCIS-tumor subgroup, which had great potential to transform to invasive cSCC.

Among Basal-SCCIS-tumor subgroup specific genes, MAGEA4 was confirmed to be strongly positive in most SCCIS and invasive cSCC by immunohistochemistry. These results suggested that MAGEA4 may play a potential biomarker of new subtype in SCCIS with more possibility into cSCC. In addition, ITGA6 and other tumor-related genes were also significantly overexpressed in Basal-SCCIS-tumor. They may play an important role in the progression of SCCIS to cSCC by regulating cell stemness, cell proliferation, cytoskeleton and extracellular matrix degradation. We also identified a group of closely related significantly up-regulated genes in cSCC. The function of genes that have received little attention in the past was validated at the cellular level. Our functional experiments found that LGALS1, IFITM3, FTH1, BST2 and SAT1 genes affected the proliferation, apoptosis, migration and invasion of human cSCC.

In TME analysis of poorly-differentiated cSCC sample, we identified major subpopulations of T lymphocytes, CAFs and DCs based on their specific markers. In cell communication analysis, we found that the vascular_Fib subpopulation with high APOE expression released high signal intensity and had the most active interaction with immune cells, especially CD8+ effector T cells. APOE gene encodes apolipoprotein, which is not only involved in lipid transport, storage and utilization, but also closely related to oxidative stress, inflammation and aging. Solé-Boldo L et al. found that the functional enrichment of these cells mainly focused on inflammatory response, wound healing, cell chemotaxis or adhesion, angiogenesis, negative regulation of cell proliferation, etc. [79]. In six DC subpopulations in cSCC, cDC1 has the most active interaction with other cells and is the main participant in the occurrence and development of cSCC. In cell-cell communication analysis, we observed significantly enhanced cell-to-cell interactions in cSCC tumor sample. The increased interactions mainly enriched in Basal1 and Pro KCs with cells in TME. In addition, we identified several crucial cancer-related signaling pathways in cSCC. The activation of these signaling pathways may play important regulatory function in cSCC tumor genesis and development.

In this study, we performed comprehensive analysis of scRNA-seq profiles in diverse samples to simulate the classic carcinogenic process from photoaged skin to AK, then to SCCIS, and finally to invasive cSCC. Especially we deeply analyzed the AK as precancerous lesions and the SCCIS at the single-cell level and identified the key malignant cell subpopulation, which is significantly important to investigate the transformation from AK to cSCC. The results are significantly benefited to understand the occurrence and development of cSCC.

## Materials and methods

### cSCC and AK patient samples

cSCCs, AKs and patient-matched normal adjacent skin samples were collected during surgical treatment at the Dermatology Department of the First Affiliated Hospital of Kunming Medical University (Yunnan, China). All AK and cSCC samples were derived from the UV-exposed areas from immunocompetent patients, and none of these patients had received any treatment before surgery. These fresh resected biopsies were divided into two parts: half of each sample was immediately dissociation into single cell suspension for single-cell sequencing, and another half was formalin fixed for pathological grading and immunohistochemical studies. Written informed consent for the samples was obtained under protocols was approved by the Ethics Committee of the First Affiliated Hospital of Kunming Medical University. Diagnosis of all samples was confirmed by at least two independent pathologists. Histological grades of cSCC were performed according to Broder’s grading system and the risk classification was performed according to the 2019 European Association of Dermato-Oncology (EADO) guidelines. And we divided AK lesions into three categories: AK Ⅰ, AK Ⅱ and AK Ⅲ, based on the abnormal cells in the percentage of intraepidermal neoplasia, as proposed by Röwert-Huber et al [80].

### Tissue dissociation

All fresh skin samples were gently washed in RPMI 1640 after removing crust, subcutaneous fat and necrotic tissue with surgical scissors and cutting the tissue into small pieces of 2–4 mm in a sterile tissue-culture dish. For tumor samples digestion was performed using tumor dissociation kit for human (130-095-929, MACS Miltenyi Biotec), mechanical dissociation using gentleMACS^TM^ Dissociator running the gentleMACS program h_tumor_01. In order to capture enough keratinocytes in AK samples for subsequent research, we separate the epidermal tissue from the dermis and then dissociated to single-cell suspensions by combining mechanical dissociation with enzymatic degradation of the extracellular adhesion proteins using Epidermis Dissociation Kit for Human (130-103-464, MACS Miltenyi Biotec). The dissociated cell suspension was strained with a 40μm filter (BD Falcon), and treated with Red Blood Cell Lysis Solution (130-103-183, MACS Miltenyi Biotec) and dead cell removal using the Dead Cell Removal Kit (130-090-101, MACS Miltenyi Biotec) to confirm cell viability >85% with trypan blue staining (Invitrogen). All samples were processed as per manufacturer’s instructions. Sorted cells were centrifuged and resuspended in PBS + 0.04% BSA (Gibco) to a final cell concentration of 700-1200 cells/μL as determined by hemacytometer.

### 10x scRNA-seq library preparation and sequencing

The single-cell capturing and downstream library constructions were performed using the Chromium Single Cell 3’ v3 (10x Genomics) library preparation kit according to the manufacturer’s protocol. Cellular suspensions were co-partitioned with barcoded gel beads to generate single-cell gel bead-in-emulsion (GEM) and polyadenylated transcripts were reverse-transcribed. Incubation of the GEMs produces barcoded, full-length cDNA from poly-adenylated mRNA, and amplified via PCR to generate sufficient mass for library construction. Then, the libraries were sequenced on NovaSeq6000 (Illumina).

### Nuclei isolation, 10x scATAC-seq library construction and sequencing

The isolation, washing, and counting of nuclei suspensions were performed according to the manufacturer’s protocol (10x Genomics, CG000169). Briefly, 100,000 to 1,000,000 cells were centrifuged at 300×g for 5 min at 4℃, removed the supernatant, and 100 µL chilled lysis buffer (10 mM Tris-HCl, 10 mM NaCl, 3 mM MgCl2, 0.1% Tween-20 and 1% BSA) was added and incubated for 5 min on ice. Following lysis, nuclei were resuspended in chilled Diluted Nuclei Buffer (10x Genomics; PN-2000153) at approximately 5,000–7,000 nuclei/µL based on the starting number of cells and immediately used to generate scATAC-seq libraries. scATAC-seq libraries were prepared according to manufacturer protocol of Chromium Single Cell ATAC Library Kit (10x Genomics, PN-1000087). Nuclei of cSCC cells were incubated by Tn5 transposable enzymes (10x Genomics; 2000138) for 60 min at 37 ℃ to form DNA frag ments. Then, mononuclear GEMs with special 10x barcodes were generated using a microfluidic platform (10x Genomics). Next, we collected single-cell GEMs and conducted linear amplification in a C1000 Touch Thermal cycler. Emulsions were coalesced using the Recovery Agent and cleaned up using Dynabeads. Indexed sequencing libraries were then constructed, purified and sequenced on NovaSeq6000 (Illumina).

### Single cell RNA-seq data processing

Reads were processed using the Cell Ranger pipeline (3.1.0) with default and recommended parameters. FASTQs generated from Illumina sequencing output were aligned to the human reference genome GRCh38-3.0.0 were generated for each individual sample by counting unique molecular identifiers (UMIs) and filtering non-cell associated barcodes. Finally, we generated a gene-barcode matrix containing the barcoded cells and gene expression counts. This output was then imported into the Seurat (4.0.5) R toolkit for quality control and downstream analysis of our single cell RNA-seq data. All functions were run with default parameters, unless specified otherwise. Low quality cells (< 200 genes/cell, < 3 cells/gene and > 10% mitochondrial genes) were excluded. Before incorporating a sample into our merged dataset, we individually inspected the cells-by-genes matrix of each as a Seurat object.

### Single cell ATAC-seq data processing

The chromatin accessibly analysis of scATAC-seq data referred to the pipeline by Muto et al [81]. The gene activity matrix was log-normalized prior to label transfer with the aggregated scRNA-seq Seurat object using canonical correlation analysis. Differential chromatin accessibility between cell types was assessed with the Signac (1.4.0) “FindMarkers” function. Genomic regions containing scATAC-seq peaks were annotated with ChIPSeeker (1.26.2) and clusterProfiler (3.18.1) using the UCSC database on hg38. Transcription factor activity was estimated using chromVAR (1.12.0) The positional weight matrix was obtained from the JASPAR2018 database. Cis-coaccessibility networks were predicted using Cicero (1.8.1)

### Identification of cell types and subtypes by nonlinear dimensional reduction

The Seurat package implemented in R was applied was applied to identify major cell types. Highly variable genes were generated and used to perform PCA. Significant principal components were determined using JackStraw analysis and visualization of heatmaps focusing on PCs 1 to 20. PCs 1 to 10 were used for graph-based clustering (at res = 0.5 for samples) to identify distinct groups of cells. These groups were projected onto UMAP analysis run using previously computed principal components 1to 10. We characterized the identities of cell types of these groups based on expression of known markers: basal cells (COL17A1, KRT5, KRT14), spinous cells (KRT1, KRT10), granular cells (FLG, LOR), proliferating keratinocytes (Pro KCs, MKI67, TOP2A), follicular cells (KRT6B, KRT17, SFRP1), Langerhans cells (CD207, CD1A), T cells (CD3D, PTPRC), melanocytes (PMEL, TYRP1) and fibroblasts (DCN, COL1A1). Sub-clustering of basal cells was further performed with the same approach.

### Cluster markers identification

The cluster-specific marker genes were identified by running the FindConservedMarkers function in the Seurat package to the normalized gene expression data. The differentially expressed genes (DEG) were identified by the ‘find.markers’ function with default parameters and filtered by p_val_adj < 0.05. Just in DEG analysis of Basal-SCCIS-tumor, we used the parameter avg_log2FC > 0.58 and p_val_adj < 0.05 to further narrow down the gene sets. We used Metascape (http://metascape.org) to perform biological process enrichment analysis with the differentially expressed genes in each cluster or subpopulation.

### CNV estimation

Initial CNVs for each region were estimated by inferCNV (1.6.0) R package. The CNV of total cell types were calculated by expression level from single-cell sequencing data for each cell with -cutoff 0.1 and -noise_filter 0.1. In order to well study the CNV level in keratinocytes for each tumor sample, we used the keratinocytes from patient-matched normal adjacent skin as background.

### Constructing single cell trajectories in keratinocytes

The Monocle3 package (1.0.0) was used to analyze single cell trajectories in order to discover the cell-state transitions. The UMI matrix was as input and variable genes obtained from epidermal cell types were detected by Seurat to sort cells in pseudotime. The subcluster of basal cells in SCCIS with higher-expressed stem cell marker (COL17A1, TP63) was defined as root stated argument and aligned via the “ordercells” function. ‘UMAP’ was applied to reduce dimensions and the visualization functions “plot_cell_trajectory’” were used to plot pseudotime trajectory.

### Haematoxylin and Eosin (H&E), Immunohistochemistry (IHC) and immunofluorescence (IF) staining

For H&E, formalin-fixed, paraffin-embedded cSCCs, AKs and patient-matched normal adjacent skin biopsies was cut at 4 microns and stained using hematoxylin and eosin (H&E). For immunohistochemistry and IF staining was performed using DAB or DAPI and the following primary antibodies: anti-ALDH3A1 mouse monoclonal antibody (Santa Cruz Biotechnology), anti-BST2 rabbit polyclonal antibody (Proteintech), anti-FTH1 rabbit polyclonal antibody (Zen-Bioscience), anti-LGALS1 rabbit polyclonal antibody (Zen-Bioscience), anti-MAGEA4 rabbit monoclonal antibody (Cell Signaling Technology), anti-IFITM3 rabbit polyclonal antibody (Zen-Bioscience), anti-IGFBP2 rabbit monoclonal antibody (Abcam), anti-ITGA6 rabbit polyclonal antibody (Zen-Bioscience), anti-SAT1 rabbit polyclonal antibody (Bioss). Examination and photographic documentation were performed using a digital slide scanner-PANNORAMIC 1000 (3DHISTECH, Hungary). Histological sections were analyzed semiquantitatively. The staining intensity of immunohistochemistry section was scored as 0 (negative), 1 (weak), 2 (medium) or 3 (strong). Extent of staining was scored as 0 (<5%), 1(5–25%), 2(26–50%), 3 (51–75%) and 4 (>75 %) according to the percentages of the positive staining areas in relation to the whole carcinoma area. Scores for staining intensity and percentage positivity of cells were then multiplied to generate the immunoreactivity score (IRS) for each case. The immunofluorescence staining was analyzed with Image-ProPlus software 6.0. Integrated option density (IOD) of interesting area (AOI) was measured and density mean (IOD/AOI) were calculated as the semi-quantitative parameters.

### Cell culture, transfections

Human immortalized epidermal keratinocytes cell line (HaCaT cells) and human cSCC cell line A431, SCL-Ⅰ and SCL-Ⅱ [82, 83] were used in this study and were obtained from the American Type Culture Collection (ATCC) and Free University of Berlin. Mycoplasma detection was carried out in cell lines using GMyc-PCR Mycoplasma Test Kit (YEASEN, 40601ES20) to avoid mycoplasma contamination. The cells were cultured in Dulbecco’s modified Eagle’s medium (DMEM, Gibico, USA) supplemented with 10% fetal bovine serum (FBS, Gibico), and 1% penicillium-streptomycin (Gibico) in a 5% CO^2^ incubator at 37°C. Small interfering RNA (siRNA) transfections were carried out using transfections reagent (INVI DNA RNA Transfections Reagent, Invigentech, USA) to inquire into the influence of silencing gene expression by on cell growth, proliferation, invasion and metastasis. The sequences of various siRNA oligonucleotides used in this study were listed in Table S20. The transfection efficiency was confirmed by RT-PCR.

### Quantitative Real-Time PCR (qRT-PCR)

To verify the expression and siRNA transfection efficiency of key genes in cSCC cells, the mRNA expression levels of genes in A431, SCL-Ⅰ and SCL-Ⅱ cells were detected by qRT-PCR. Primers were designed and listed in Table S21. Total RNA was extracted from cells using TRIzol reagent (Invitrogen, Thermo Fisher Scientific, USA) and reverse transcribed into cDNA using a FastKing-RT Reagent kit (Tiangen, Beijing, China) according to the manufacturer’s protocols. Quantitative RT-PCR was performed using SYBR Green Master Mix (Tiangen, Beijing, China). The RNA expression level of target genes was evaluated _by 2_−ΔΔCt.

### Cell-counting kit 8(CCK8) assay

CCK8 assay was employed for the evaluation of cell proliferation. The transfected cells were seeded in a 96-well plate at a seeding density of 2000 cells/well (100μL). Next,10 μL CCK8 reagent (Beyotime, Shanghai, China) was added to each well and incubated at 37℃. Cell proliferation rate was assessed according to the optical density (OD) value (450 nm) detected by Microplate reader (BioTek, USA) at 0 h, 24 h, 48 h and 72 h following the instructions of manufacture.

### Annexin V and PI staining detects cell apoptosis

The apoptosis rate was evaluated using the Annexin V-FITC and propidium iodide (PI) kit (Beyotime, Shanghai, China) following the manufacturer’s protocols. A431, SCL-Ⅰ and SCL-Ⅱ cells were transfected and suspended, and 5 μl Annexin V-FITC and 10 μl PI were added to the cell suspension. After 20 minutes of incubation at room temperature in the dark, the cells were analyzed by flow cytometry (FACS Cabibur; BD, CA, USA).

### Cell migration assay

A wound-healing assay was performed to test the migration ability of the cSCC cells after transfection. Cells were grown to confluence in 6-well plates, and the wounds were made in confluent monolayer cells using a sterilized 200 µL pipette tips. The culture medium was then removed, and the cells were washed with PBS and cultured with the indicated treatment. Wound healing of different groups was detected at 0, 24, 48, and 72 hours within the scraped lines, and representative fields were photographed at the different time points to assess the migratory ability of the cells.

### Transwell invasion assays

The invasion ability of cSCC cells was evaluated by Transwell assays using Transwell chambers (8 μm pore size, Corning Costar, USA) precoated with Matrigel. After transfection, 4×10^4^ cSCC cells in the 200 μL serum-free medium were added to the upper chambers, DMEM with 10% FBS was added to the bottom chambers. After incubation at 37°C for 36 h, the cells invaded into the lower side of the inserts were fixed in 4% paraformaldehyde and stained with 0.1% crystal violet. Then counted and photographed under a microscope.

### TME analysis

On the basis of the general cell population, we extracted T cells、DC cells and stromal cells individually for further subdivision. For subgroup cell clustering, cells of different types were extracted separately and clustered by their respective parameters (T cells: 21 PCs, using resolution of 0.5 (CD8_T: 18 PCs, using resolution of 0.7; CD4_T: 20 PCs, using resolution of 1); DC: 24 PCs, using resolution of 0.9; Fib: 26 PCs, using resolution of 0.3). The annotations of cell identity on each subcluster were defined by the expression of known marker genes. Inference of intercellular communications was conducted using ‘CellChat’. The ligand-receptor interactions were all based on ‘CellChatDB’, a database of literature-supported ligand-receptor interactions in both mouse and human. The majority of ligand–receptor interactions in CellChatDB were manually curated on the basis of KEGG (Kyoto Encyclopedia of Genes and Genomes) signaling pathway database. The identification of major signals for specific cell groups and global communication patterns was based on an unsupervised learning method non-negative matrix factorization.

### Statistical analysis

All experiments were performed in triplicate technical replicates, and all data are presented as mean ± standard deviation (SD). Differences among groups were analyzed using Student’s t test or one-way analysis of variance (ANOVA) for normally distributed data and the Kruskal-Wallis test for non-normally distributed data. And P < 0.05 was considered to be statistically significant.

## Acknowledgments

The authors acknowledge the editors and reviewers for their positive and constructive comments and suggestions related to this study.

## Author contributions

L.H. and X.L. conceived and designed the study. D.Z., J.Q., Y.T., X.L., F.L., L.H., and H.L. procured and processed tissue specimens. D.Z. prepared single-cell suspensions, sequencing libraries and performed next-generation sequencing. Y.S., X.L., Y.H. performed quality checks, data integration, and computational analyses. Y.S., D.Z., X.L., L.H., X.L., W.W. and D.X. analyzed and interpreted scRNA-seq data. D.Z. and X.W. performed the IHC, IF staining and cytology experiments. Y.S., D.Z., X.L., X.L., L.H., W.W. and D.X. wrote and revised manuscript. All authors reviewed the results and approved the final version of the manuscript.

## Availability of data and materials

The raw data and gene counts table are available from GEO under accession number (GSE193304). All data needed to evaluate the conclusions in the paper are present in the paper and/or the Supplementary Materials. Additional data related to this paper may be requested from the authors.

## Ethical approval and consent to participate

The authors are accountable for all aspects of the work in ensuring that questions related to the accuracy or integrity of any part of the work are appropriately investigated and resolved. All procedures performed in this study involving human participants were in accordance with the *Declaration of Helsinki* (as revised in 2013). This study protocol was approved by the Ethics Committee of the First Affiliated Hospital of Kunming Medical University (Approval Number (2020)-L-29), and written informed consent was obtained from all patients.

## Supplementary Figures

**Fig. S1.**
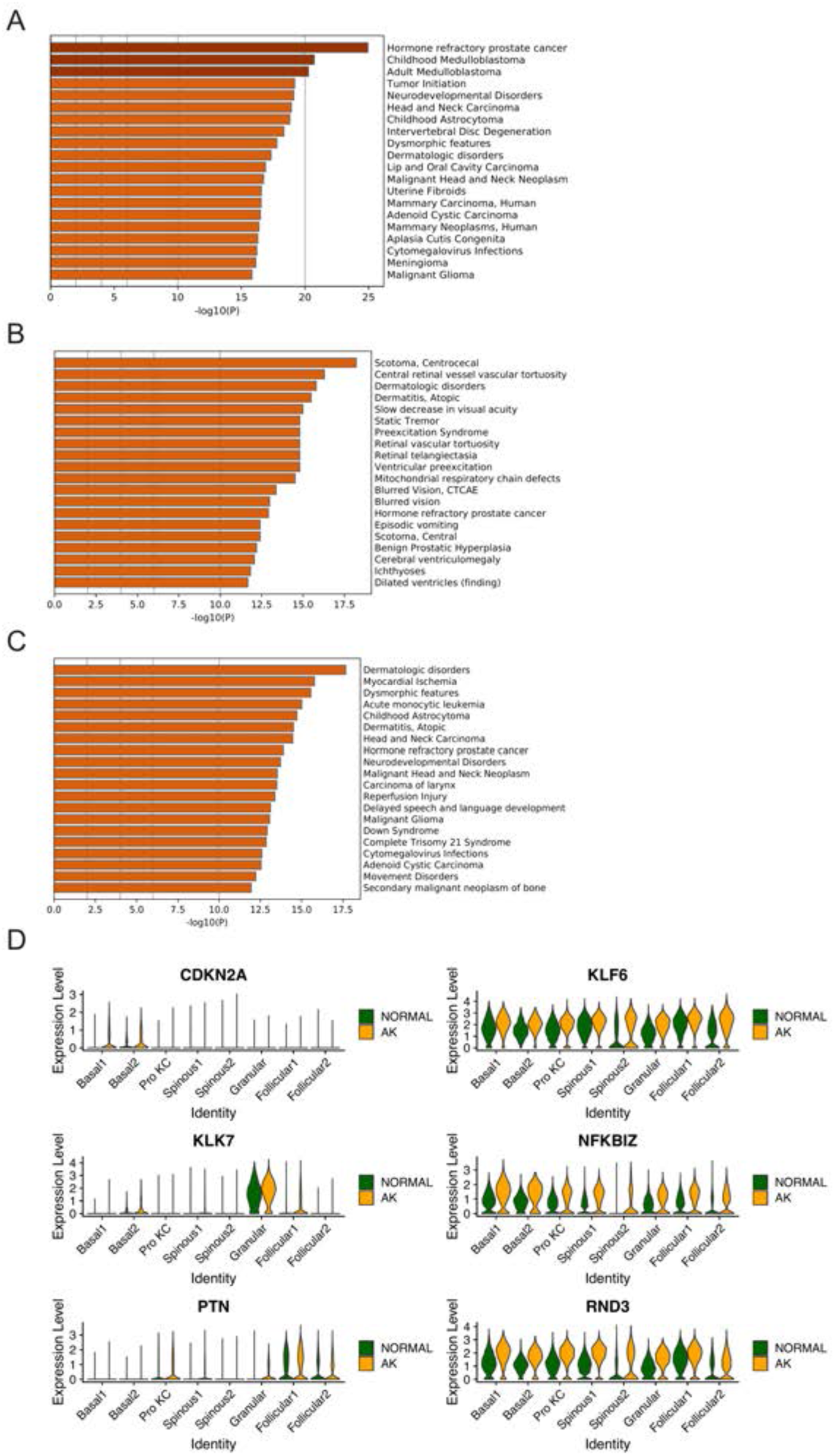
Identification of potential key driver genes from normal skin to AK. (A) The enriched disease terms of up-regulated DEGs in Basal1 between AK and normal groups based on DisGeNET database. (B) The enriched disease terms of up-regulated DEGs in Basal2 between AK and normal groups based on DisGeNET database. (C) The enriched disease terms of up-regulated DEGs in Pro KC between AK and normal groups based on DisGeNET database. (D) Violin plots showing the different expression levels of candidate genes in AK and normal samples.

**Fig. S2.**
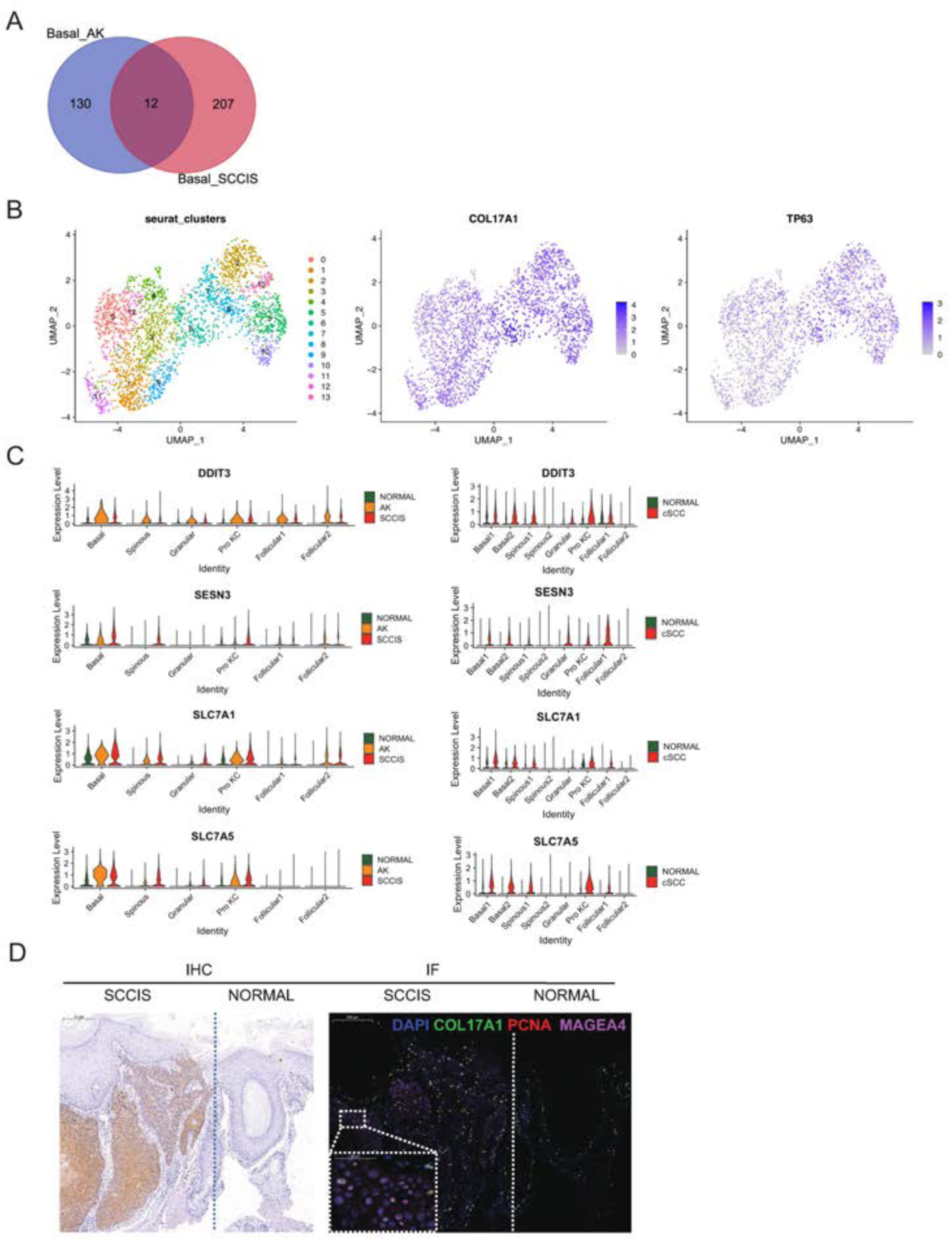
The comprehensive analysis of the patient (P2) with both AK and SCCIS. (A) Overlap of down-regulated genes in basal cells from AK compared to normal and SCCIS compared to AK. (B) Left, UMAP of subgroups generated from basal cells in SCCIS sample labeled by Seurat clusters; middle and right, expression of stem cell marker (COL17A1, TP63) in basal cells in SCCIS sample. (C) Violin plots showing the different expression levels of candidate genes across all types of keratinocytes in P2, cSCC and normal groups. (D) Left, immunohistochemical staining showed the expression of MAGEA4 in SCCIS (left) was higher than that for para-cancer normal skin tissues (right). Scale bar, 200 μm. Right, immunofluorescence staining for COL17A1 (green), PCNA (red) and MAGEA4 (pink) validates their co-expression in SCCIS. DAPI stains nuclei. Scale bar, 200 μm. The representative views of co-staining were shown in the enlarged images at bottom left. Scale bar, 50 μm.

**Fig. S3.**
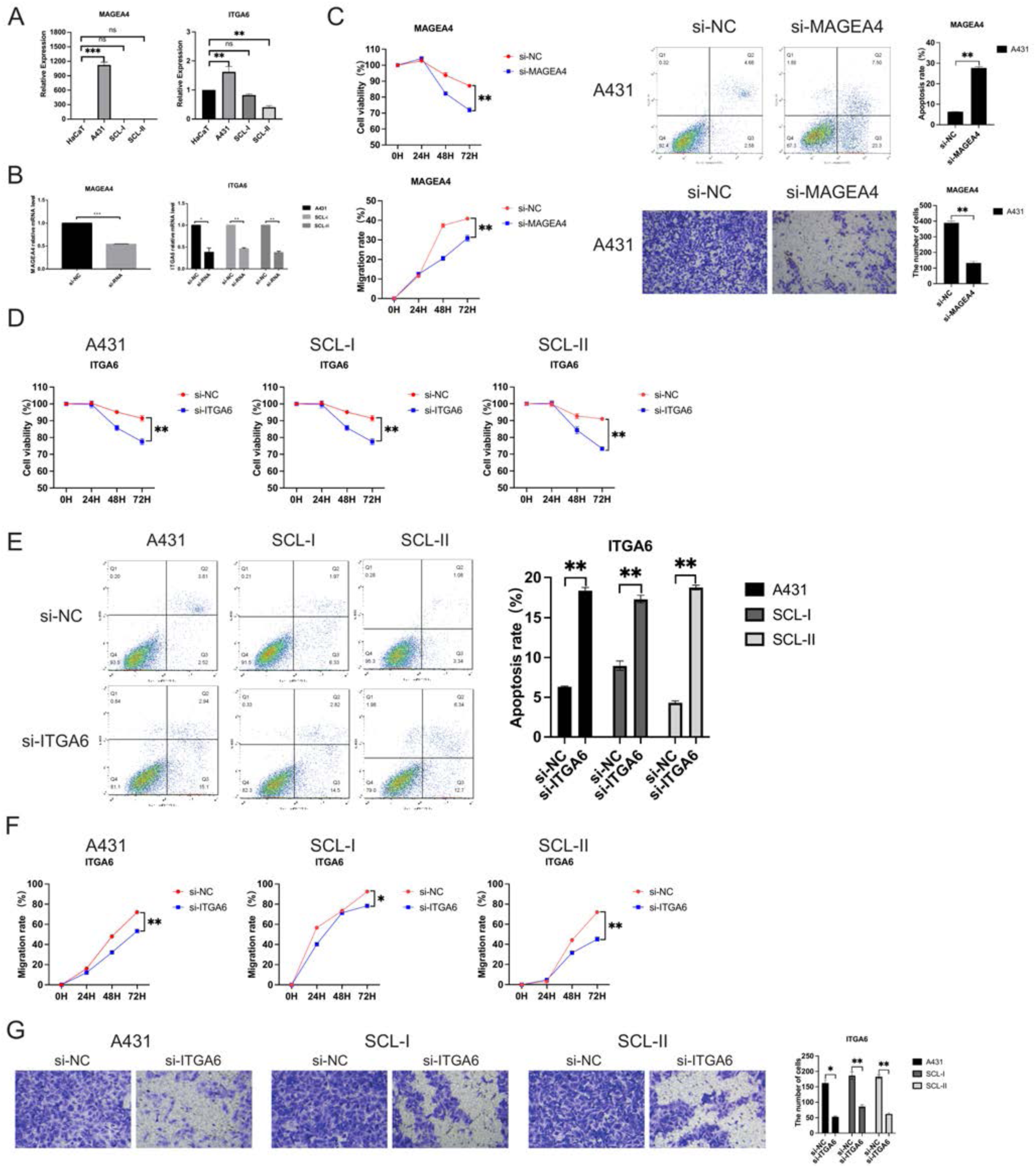
Expression and functional characterization of MAGEA4 and ITGA6. (A) The mRNA expression of MAGEA4 and ITGA6 in human immortalized keratinocytes (HaCaT) and cSCC cell lines (A431, SCL-I, SCL-II). *p < 0.05; **p < 0.01; ***p < 0.001; ns, not significant. (B) Effect of siRNA on the expression of MAGEA4 in A432 and ITGA6 in A431, SCL-I and SCL-II determined by qRT-PCR. (C) Functional experiment of MAGEA4 in A432. Upper, left, effect of MAGEA4 cSCC cell proliferation by CCK-8 proliferation in A431; upper, right, the effect of MAGEA4 on cSCC cell apoptosis was measured by staining with Annexin V-FITC/PI, followed by FACS analysis. Lower, left, the scratch experiment showed that MAGEA4 knockdown resulted in a shorter vertical migration distance compared with the control group after 72 h; lower, right, transwell assay showed that the invasion abilities of the si-MAGEA4 groups significant decreased compared with the si-NC group. **p < 0.01. (D) Effect of ITGA6 on cSCC cell proliferation by CCK-8 proliferation assay in A431, SCL-I and SCL-II. *p < 0.05; **p < 0.01. (E) The effect of ITGA6 on cSCC cell apoptosis. **p < 0.01. (F) ITGA6 knockdown resulted in a shorter vertical migration distance compared with the control group after 72 h. *p < 0.05; **p < 0.01. (G) The invasion abilities of the si-ITGA6 groups significant decreased compared with the si-NC group. *p < 0.05; **p < 0.01.

**Fig. S4.**
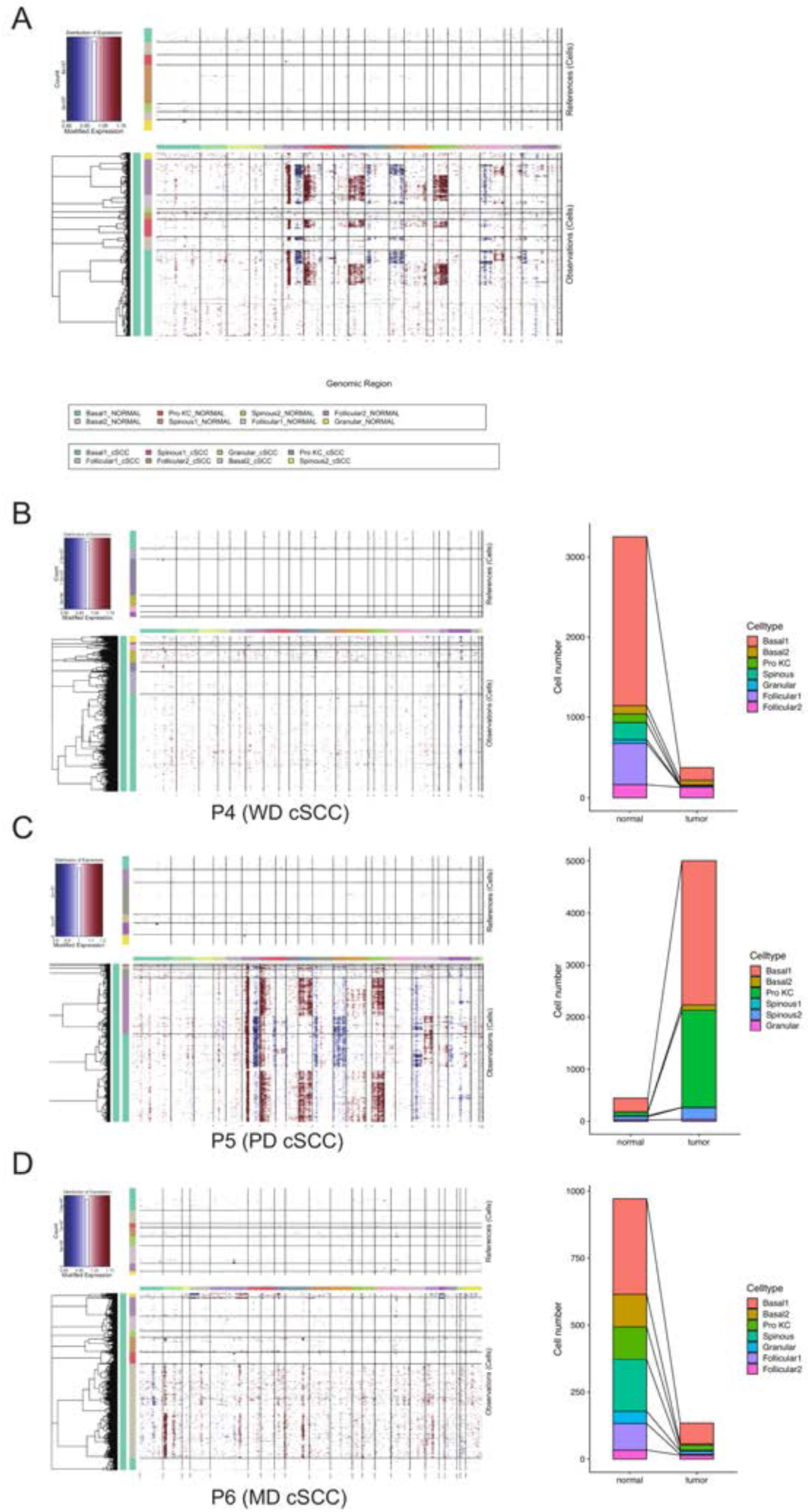
CNV scores positively correlated with malignant degrees of cSCC. (A) Heatmap showing CNV levels of all keratinocytes from all cSCC samples. The keratinocytes from all patient-matched normal samples were defined as references. (B) Left, heatmap showing CNV levels of all keratinocytes from WD cSCC sample; right, the proportion of tumor and normal cells in WD cSCC sample defined by cnv.cut (probs = 0.99). (C) Left, heatmap showing CNV levels of all keratinocytes from PD cSCC sample; right, the proportion of tumor and normal cells in PD cSCC sample defined by cnv.cut (probs = 0.99). (D) Left, heatmap showing CNV levels of all keratinocytes from MD cSCC sample; right, the proportion of tumor and normal cells in MD cSCC sample defined by cnv.cut (probs = 0.99).

**Fig. S5.**
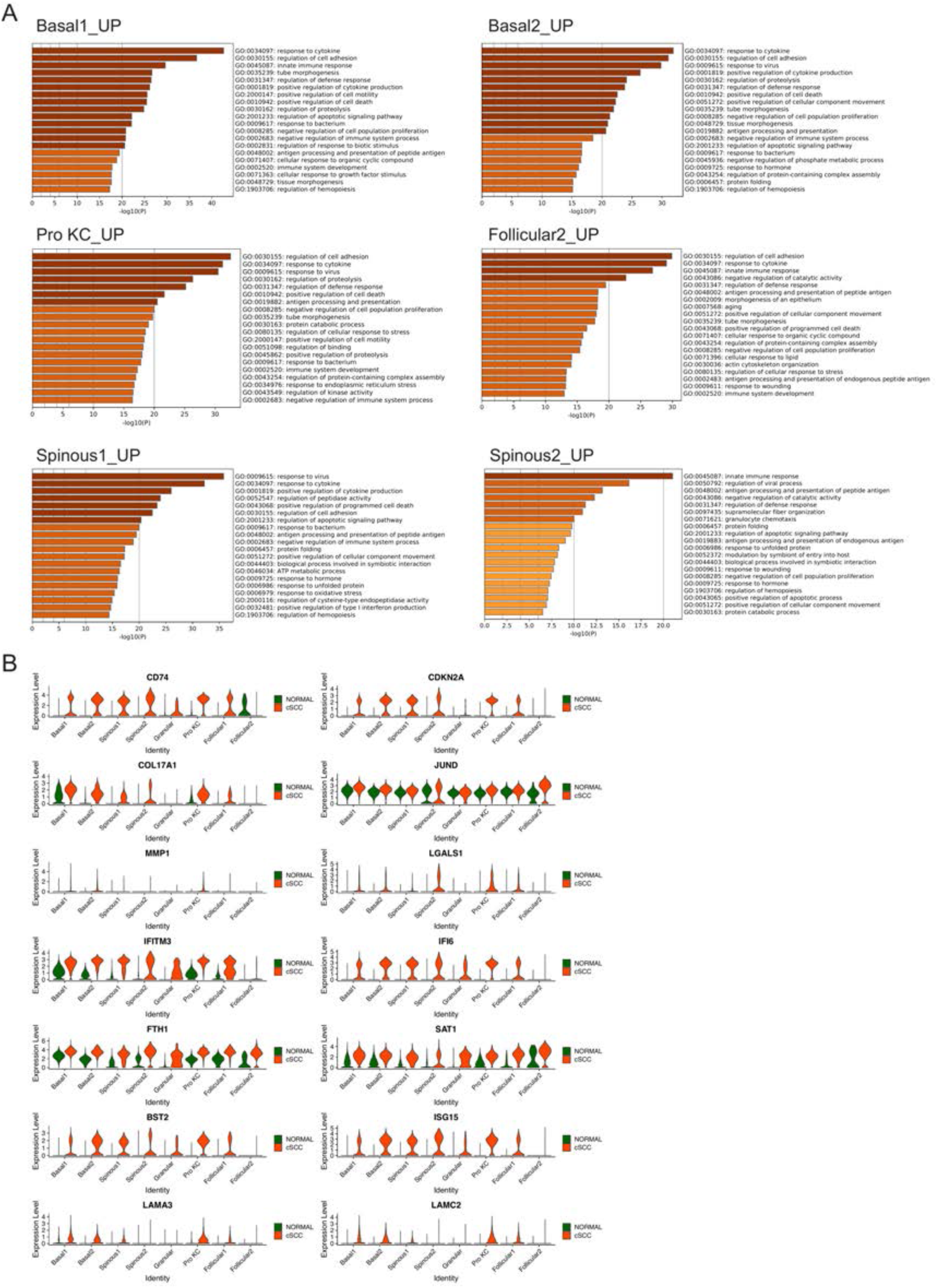
Identification of key genes associated with cSCC. (A) The enriched GO terms of up-regulated DEGs in Basal1, Basal2, Pro KC, Follicular2, Spinous1and Spinous2 between cSCC and normal groups. (B) Violin plots showing the different expression levels of candidate genes in cSCC and normal groups.

**Fig. S6.**
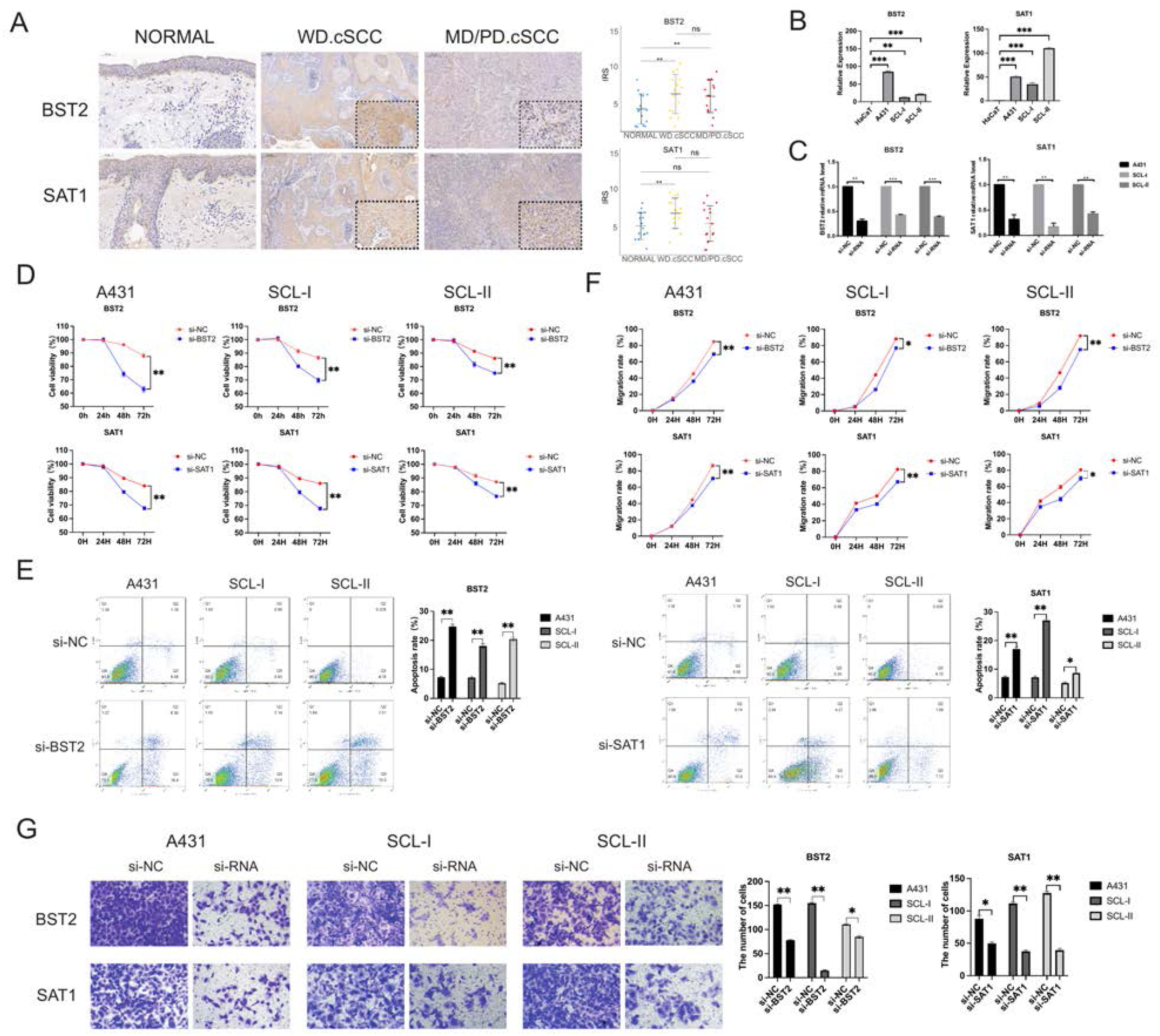
Expression and functional characterization of BST2 and SAT1. (A) Left, immunohistochemical staining of BST2 and SAT1 in cSCC in normal skin (200X), WD cSCC (50X & 250X) and MD/PD cSCC (50X & 250X). Scale bar, 200 μm & 50 μm. Right, The immunoreactivity score (IRS) analyses of BST2 and SAT1 in normal skin, WD cSCC and MD/PD cSCC. n = 15 for each group. *p < 0.05; **p < 0.01; ***p < 0.001; ns, not significant. (B) The mRNA expression of BST2 and SAT1 in human immortalized keratinocytes (HaCaT) and cSCC cell lines (A431, SCL-I, SCL-II). *p < 0.05; **p < 0.01; ***p < 0.001; ns, not significant. (C) Effect of siRNA on the expression of BST2 and SAT1 in A431, SCL-I and SCL-II determined by qRT-PCR. (D) Effect of BST2 and SAT1 on cSCC cell proliferation by CCK-8 proliferation assay in A431, SCL-I and SCL-II. **p < 0.01. (E) The effect of BST2 and SAT1 on cSCC cell apoptosis. **p < 0.01. (F) BST2 and SAT1 knockdown resulted in a shorter vertical migration distance compared with the control group after 72 h. *p < 0.05; **p < 0.01. (G) The invasion abilities of the si-ITGA6 groups significant decreased compared with the si-NC group. *p < 0.05; **p < 0.01.

**Fig. S7.**
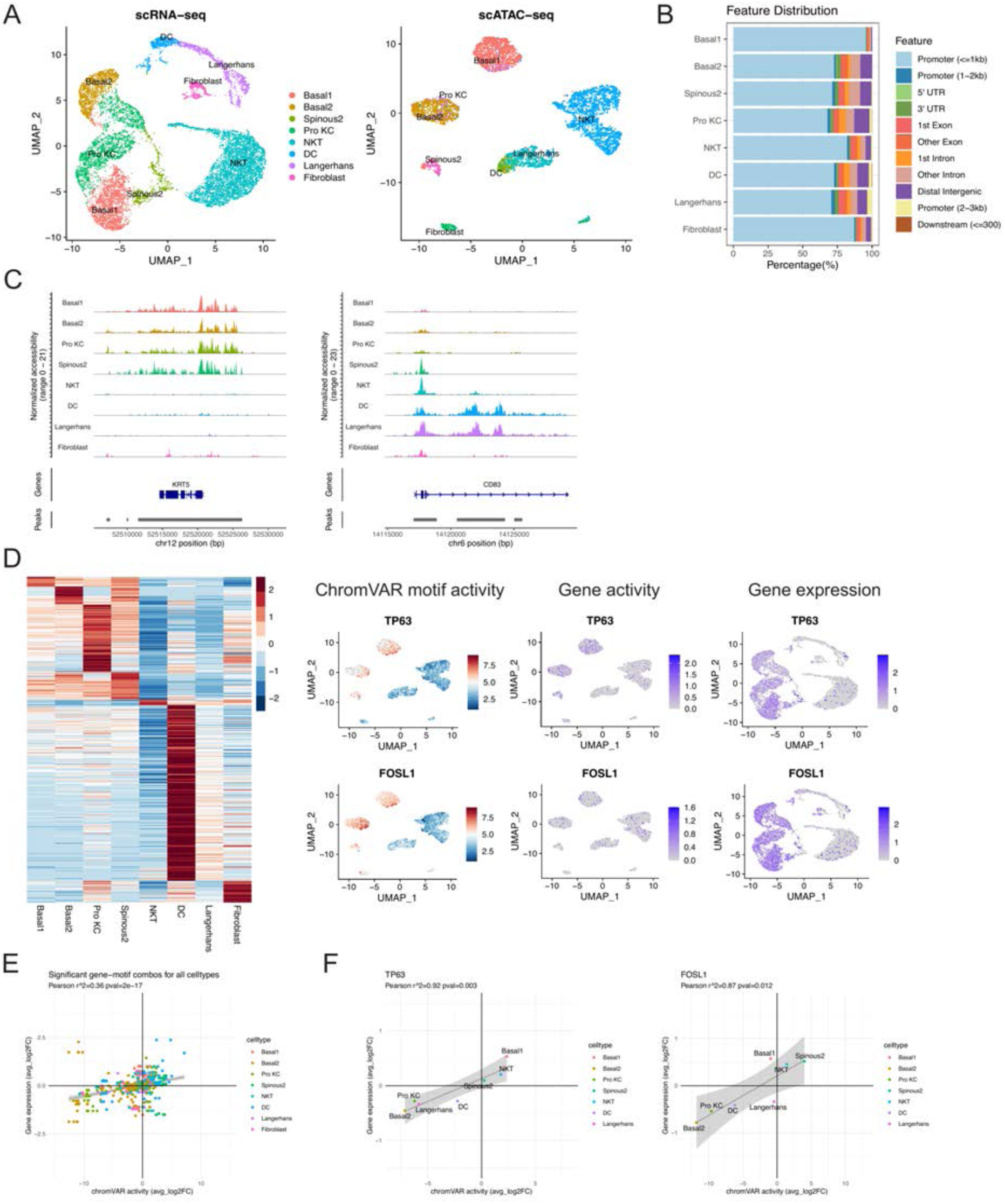
Chromatin accessibility is associated with transcription factor activity. (A) Left, UMAP of scRNA-seq dataset from PD cSCC sample labeled by cell type; right, UMAP plot of scATAC-seq dataset from PD cSCC sample labeled by cell type after integration and label transfer with scRNA-seq data. (B) Bar plot of annotated differentially accessible region (DAR) location for each type. (C) Fragment coverage (frequency of Tn5 insertion) around the DAR on the gene KRT5 and CD83. (D) Left, Heatmap of average chromVAR motif activity for each cell type. The color scale represents a z-score scaled by row. Right, UMAP plot displaying chromVAR motif activity, gene activity and gene expression of TP63 (upper) and FOSL1 (lower). The color scale for each plot represents a normalized log-fold-change for the respectively assay. (E) Cell-specific mean chromVAR motif activity from the JASPAR database was plotted against cell-specific average expression for the corresponding transcription factor for all cell types and transcription factors. (F) Mean chromVAR activity was plotted against average expression for TP63 (left) and FOSL1 (right). Significant correlation was assessed with Pearson’s product moment correlation coefficient using the cor.test function in R.

## See zip file

**Table S1-S16 S19-S21.**

**Table S1:** Clinical characteristic of patients and samples enrolled in single-cell sequencing. **Table S2:** The gene list of up-regulated DEGs in AK Basal1 subpopulation. **Table S3:** The gene list of up-regulated DEGs in AK Basal2 subpopulation. **Table S4:** The gene list of up-regulated DEGs in AK Pro KC subpopulation. **Table S5:** AK candidate driver genes and antibodies for IF. **Table S6:** The gene list of overlapped up-regulated DEGs in Basal subpopulation of P2 from AK vs normal and SCCIS vs AK.. **Table S7:** The gene list of overlapped down-regulated DEGs in Basal subpopulation of P2 from AK vs normal and SCCIS vs AK. **Table S8:** The gene list of up-regulated DEGs between Basal-SCCIS-tumor vs Basal-SCCIS-normal. **Table S9:** SCCIS candidate driver genes and antibodies for IHC. **Table S10:** The gene list of up-regulated DEGs in cSCC Basal1 subpopulation. **Table S11:** The gene list of up-regulated DEGs in cSCC Basal2 subpopulation. **Table S12:** The gene list of up-regulated DEGs in cSCC Pro KC subpopulation. **Table S13:** The gene list of up-regulated DEGs in cSCC Follicular2 subpopulation. **Table S14:** The gene list of up-regulated DEGs in cSCC Spinous1 subpopulation. **Table S15:** The gene list of up-regulated DEGs in cSCC Spinous2 subpopulation. **Table S16:** cSCC candidate driver genes and antibodies for IHC. **Table S19:** The correlation between chromVAR transcription factor activity with expression in scATAC-seq data. **Table S20:** The sequences of various siRNA oligonucleotides used in this study. Table 21: The primers of genes used for qRT-PCR.

**Table S17.** The differentially accessible chromatin regions between cell types of poorly-differentiated cSCC sample in scATAC-seq data.

**Table S18.** The chromVAR transcription factor activity between cell types of poorly-differentiated cSCC sample in scATAC-seq data.

